# Hedgehog regulation of epithelial cell state and morphogenesis in the larynx

**DOI:** 10.1101/2022.01.19.476887

**Authors:** Janani Ramachandran, Weiqiang Zhou, Anna E. Bardenhagen, Talia Nasr, Aaron M. Zorn, Hongkai Ji, Steven A. Vokes

## Abstract

The larynx enables speech while regulating swallowing and respiration. Larynx function hinges on the laryngeal epithelium which originates as part of the anterior foregut and undergoes extensive remodeling to separate from the esophagus and form vocal folds that interface with the adjacent trachea. Here we find that Sonic hedgehog (SHH) is essential for epithelial integrity in the larynx as well as the anterior foregut. During larynx-esophageal separation, low *Shh* expression marks specific domains of actively remodeling epithelium that undergo an epithelial to mesenchymal transition (EMT) characterized by the induction of N-Cadherin and movement of cells out of the epithelial layer. Consistent with a role for SHH signaling in regulating this process, *Shh* mutants undergo an abnormal EMT throughout the anterior foregut and larynx, marked by a cadherin switch, movement out of the epithelial layer and cell death. Unexpectedly, *Shh* mutant epithelial cells are replaced by a new population of *Pax-1* expressing cells that form a rudimentary epithelium. These findings have important implications for interpreting the etiology of HH-dependent birth defects within the foregut. We propose that SHH signaling has a default role in maintaining epithelial identity throughout the anterior foregut and that regionalized reductions in SHH trigger epithelial remodeling.

## Introduction

The larynx produces all of the sounds for vocal communication and regulates swallowing and access to the esophagus and trachea that lie directly beneath it. Congenital laryngeal malformations such as tracheo-laryngeal clefts and bifid epiglottis arise from defects in early laryngeal morphogenesis and impair trachea-esophageal function (feeding and breathing), as well as vocalization in infants, often requiring surgical intervention and significantly impacting patients’ quality of life (Biesecker, 1993; Cohen et al., 2006; Johnston et al., 2014; Leboulanger and Garabédian, 2011). Complicating the etiology of these disorders, the pathways that drive the early stages of larynx morphogenesis, specifically vocal fold closure and larynx-esophageal separation, remain largely unknown. Several recent findings suggest that HH signaling may be important for early larynx development. Early loss of HH signaling results in the reduction of SOX2-expressing cells from the larynx epithelium and a failure in vocal fold closure (Lungova et al., 2015). HH signaling also drives the separation of the trachea and esophagus, which are directly caudal to the larynx (Billmyre et al., 2015; Han et al., 2020; Ioannides et al., 2010; Kim et al., 2019; Kuwahara et al., 2020; Nasr et al., 2019; Que et al., 2007). Mutations in the HH pathway transcriptional effector GLI3 cause dramatically altered larynx morphology and vocalization defects (Tabler et al., 2017). Similarly in humans, laryngeal clefts and bifid epiglottis are phenotypes of Pallister Hall syndrome which arises from truncating mutations in GLI3 (Biesecker, 1993; Bose, 2002; Ondrey et al., 2000). Together these defects suggest that HH signaling may be required for several stages of larynx morphogenesis beyond vocal fold closure.

The larynx is derived from the early foregut epithelium which is regionally differentiated into multiple organs, including the pharynx, parathyroid, thymus, trachea, esophagus and larynx in the anterior half. Induction of these organs from the nascent gut tube, as well as subsequent morphogenesis are driven by specialized types of epithelial remodeling such as budding, branching, septation, and epithelial to mesenchymal transitions (EMT) (Bort et al., 2006, 2004; Hebrok, 2000; Hogan and Kolodziej, 2002; Qi and Beasley, 2000), and are regulated by localized signaling interactions, including HH, between the foregut and the surrounding mesenchyme (Han et al., 2020, 2017; Jacobs et al., 2012; Kraus and Grapin-Botton, 2012; Nerurkar et al., 2017; Rankin et al., 2016; Zorn and Wells, 2009). In the anterior foregut, organogenesis is uniquely affected by the influx of migratory neural crest-derived cell populations that combine with populations of mesodermally-derived mesenchymal cells to form region-specific pharyngeal structures (Bain et al., 2016; Bhatt et al., 2013; Brito et al., 2006; Kuo and Erickson, 2010; Tabler et al., 2017; Trainor and Tam, 1995).

Arising initially from the 4^th^ pharyngeal pouch and the pharyngeal floor (Figure 1A), the larynx bridges the anterior-most portions of the foregut to the more posterior trachea and esophagus (Henick, 1993; Lungova et al., 2015). The early stages of larynx development are characterized by three major epithelial remodeling events, beginning with the stratification and zippering of the lateral walls of the foregut along the midline to close the vocal folds and form the epithelial lamina. Within the next 24 hours, the epithelial lamina, which joins the dorsal esophagus and the ventral trachea, puckers to form the infraglottic duct, and separates from the esophagus (Henick, 1993; Lungova et al., 2015). The newly separated lamina then fully recanalizes to form a laryngeal lumen that is continuous with the trachea, around which specialized cartilage elements and musculature are specified (Henick, 1993; Lungova et al., 2018, 2015). Epithelial morphogenesis is genetically dependent upon Wnt, Hippo and HH signaling although the underlying mechanisms remain poorly understood (Lungova et al., 2018, 2015; Mohad et al., 2021; Tabler et al., 2017).

**Figure 1.**
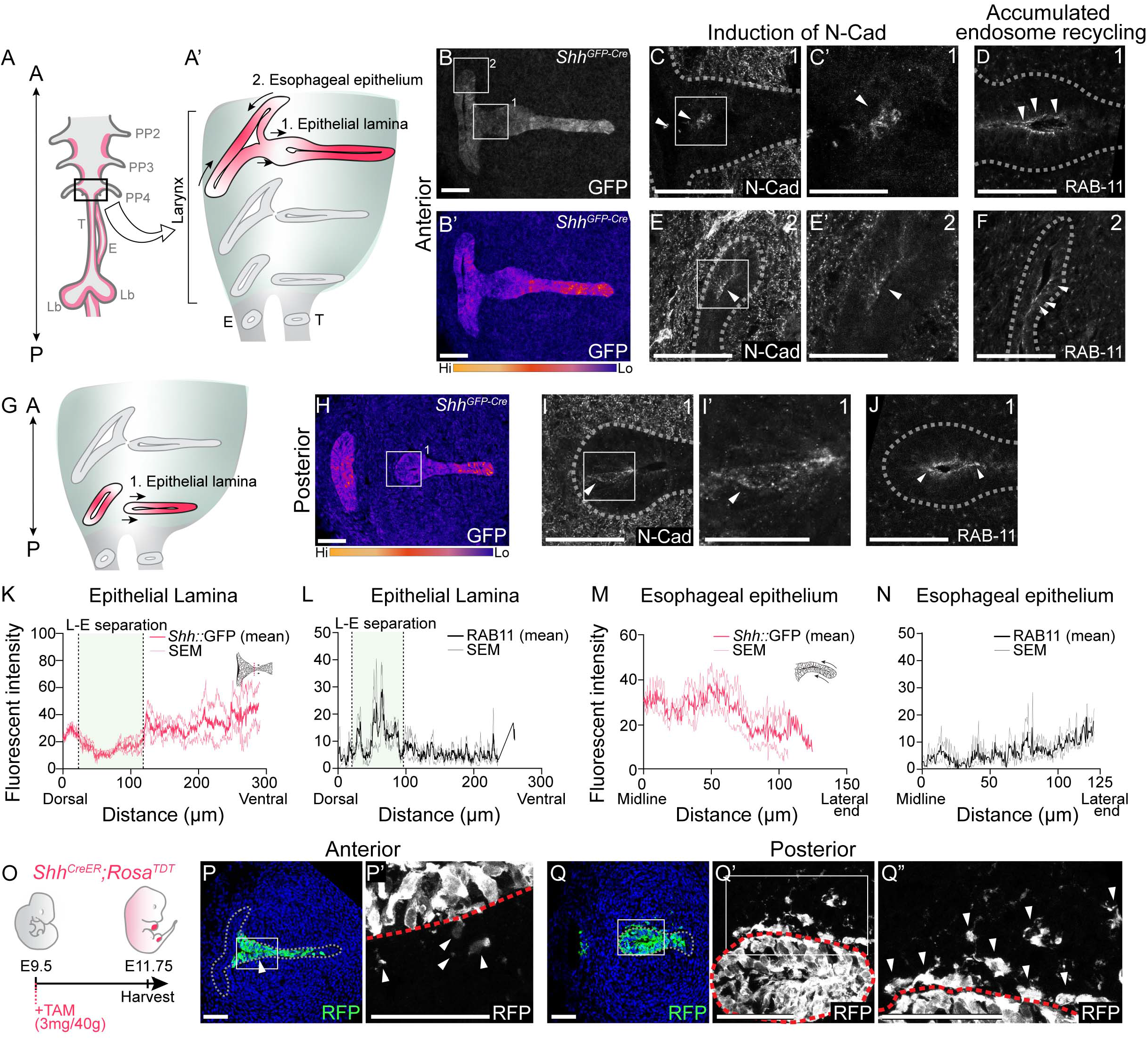
Actively remodeling epithelial cells have low *Shh* expression and undergo EMT during larynx- esophageal separation and esophageal constriction. A, G. Schematic of the anterior foregut (A) and highlight of the anterior (A) and posterior (G) larynx at E11.75. A, B, H GFP marking *Shh* expression in the anterior and posterior larynx (n=5 for each). There is reduced GFP expression at the epithelial lamina which fuses and then separates the larynx and esophagus (region 1; B,H), and the constricting esophageal opening (region 2; B) of the larynx. C,E-E’,I-I’. Expression of N-Cadherin at anterior or posterior regions at E11.75 (n=3). D,F,J. RAB-11 was visualized at regions 1 (D,J) and 2 (F) in 3 larynxes. K, M. Relative GFP expression along the epithelium at regions 1 (K) and 2 (M) was measured and averaged across 3 replicates by line scans of fluorescent intensity. Standard error of mean was calculated across all 3 replicates and plotted in light pink. L,N. Relative RAB-11 expression was measured by lines scans of fluorescent intensity along the epithelium at regions 1 (L) and 2 (N) and averaged across 3 replicates. Standard error of mean was calculated across all 3 replicates and plotted in gray. O-Q. Shh-descendant cells were visualized in 3 E11.75 larynxes using *Shh^CreER^;Rosa^TDT^* lineage labeling. *Shh^CreER/+^;Rosa^TDT/+^* embryos were induced with Tamoxifen at E9.5 and analyzed for RFP (green) expression (P-Q) at E11.75 along the anterior-posterior axis of the larynx. A-anterior; P-posterior; PP 2/3/4, Pharyngeal pouches #2-4, Lar-larynx, T-trachea, E-esophagus, Lb-lung buds. C’,E’,I’. Scale bars denote 25pm. All other scale bars denote 50pm. 30

We asked if and how HH signaling might regulate epithelial remodeling during larynx development. We defined distinct domains of epithelium that downregulate *Shh* and undergo EMT-based remodeling during larynx-esophageal separation and esophageal constriction. We uncovered a similar process in *Shh^-/^*^-^ embryos, in which epithelial cells lose expression of canonical foregut genes and undergo an EMT marked by cadherin switching and ultimately cell death. Despite massive cell death, the anterior foregut retains a rudimentary epithelium that now contains an ectopic population of cells. These findings provide a cell-based mechanism for understanding previously defined HH-dependent vocal fold closure and laryngeal cleft defects (Lungova et al., 2015). As similar changes are seen beyond the larynx, we propose a model in which regionalized reductions in HH drive dynamic epithelial remodeling throughout the anterior foregut.

## Results

### Larynx epithelial cells lose *Shh* expression and undergo EMT based remodeling during larynx-esophageal separation, and esophageal constriction

To determine if *Shh* might regulate epithelial remodeling in the larynx, we examined its expression at E11.75, when the vocal folds are remodeled to separate the larynx from the esophagus. There was a wide variation in *Shh* expression within the larynx, with markedly reduced domains of *Shh^GFP^* expression in the epithelial lamina at the future site of larynx-esophageal separation in addition to the lateral edges of the esophagus that are in the process of constricting (Figure 1A- B, G-H, K,M). The regional reduction in GFP expression is corroborated by a loss of *Shh* gene expression in both regions (Figure1-figure supplement 1A-D), as well as the absence of *Shh*- descendant cells from these regions (Figure1-figure supplement 2A-B).

Because *Shh* was reduced in both regions of the larynx undergoing dynamic epithelial remodeling, and previous studies observed *Shh*-descendant cells in the mesenchyme directly adjacent to larynx-esophageal separation (Lungova et al., 2018), we asked whether the absence of *Shh* in the larynx epithelium was accompanied by cadherin-switching and EMT. Consistent with this possibility, membranous N-Cadherin was expressed within the epithelial layer both in the epithelial lamina adjacent to the infraglottic duct at the site of larynx-esophageal separation, as well as along the lateral edges of the constricting esophagus (Figure 1C, E). N-Cadherin-expressing cells were also present in more caudal sections of the separated larynx, overlapping with the region of reduced GFP expression (Figure 1I). While there was no overall reduction in E-Cadherin protein (Figure1-figure supplement 1E-I), there was an increase in punctate E-cadherin expression in both regions (Figure1-figure supplement 1H’, I’). The re-localization of E-Cadherin and the concomitant initiation of N-Cadherin at these regions provides evidence for a Cadherin-switch both at the epithelial lamina and along the lateral edges of the esophagus. This is further supported by the apical accumulation RAB-11, a marker of endosome recycling that is required for the transport of E-Cadherin as well as N-Cadherin to the apical cell surface in multiple contexts (Desclozeaux et al., 2008; Kawauchi et al., 2010; Nasr et al., 2019; Welz et al., 2014; Woichansky et al., 2016) (Figure 1D,F,J,L,N).

To determine whether N-Cadherin expressing cells undergo EMT within these domains we examined *Shh*-descendant cells at E11.75 stage using a *Shh^CreER^;Rosa^TDT^* reporter line (Figure 1O). Consistent with prior reports using a related *Shh^Cre^* strategy (Lungova et al., 2018), there were a small number of RFP-labeled cells within the mesenchyme along the anterior-posterior axis of the separating larynx and esophagus (Figure 1P-Q). While there was a significant increase in the domain of N-Cadherin expression within the remodeling epithelium at later stages of larynx- esophageal separation (Figure 1-figure supplement 2D-F), there was no significant increase in the number of mesenchymal RFP-labeled cells (Figure 1-figure supplement 2B-C). This suggests that some *Shh*-descendant cells undergo EMT-based extrusion during larynx remodeling but they do not remain in the mesenchyme. Overall, these findings indicate that *Shh* expression is dynamically regulated in the remodeling larynx, with low levels of *Shh* expression coinciding with cadherin switching. The change in cadherin status is likely the underlying cause for the epithelial cells to leave the epithelium by an EMT-like process.

### Larynx epithelial cells undergo ectopic EMT-like cell extrusion in the absence of HH signaling

Our results so far indicated that regional loss of *Shh* was associated with cadherin switching and EMT. To investigate this further, we generated RNA-seq datasets for control and *Shh^-/-^* larynx tissues and identified differentially regulated genes and enriched pathways (Supplemental Data Table 1). Consistent with this model, epithelial to mesenchymal transition (EMT) was the most significantly enriched pathway among HH-dependent genes (Figure 2A), supporting a role for HH signaling in regulating this process. Differentially expressed genes consisted of members of all three progressive EMT stages (Figure 2B) (Lamouille et al., 2014). These included downregulation of the pro-epithelial adhesion genes *Dsp*, and *Dcn,* which mark the first stage (Bax et al., 2011; Huang et al., 2012; Kowalczyk and Nanes, 2012; Wang et al., 2015; Yilmaz and Christofori, 2009). There was also an upregulation of the pro-migratory genes *Cdh2, Vimentin,* and *Fn1,* indicative of the next phase of EMT (Wheelock et al., 2008; Yilmaz and Christofori, 2009). Finally, there was a downregulation of *Lama1*, suggestive of a breakdown in basement membrane which is one indicator of the third stage of EMT (Aumailley and Smyth, 1998; Lamouille et al., 2014; Nakaya et al., 2008; Thiery and Chopin, 1999). In addition, several TGFβ family members were up-regulated, suggesting a plausible mechanism for inducing EMT (Barrallo-Gimeno and Nieto, 2005; Katsuno et al., 2013; Mercado-Pimentel and Runyan, 2007; Nawshad et al., 2004; Schnaper et al., 2003; Thiery et al., 2009).

**Figure 2.**
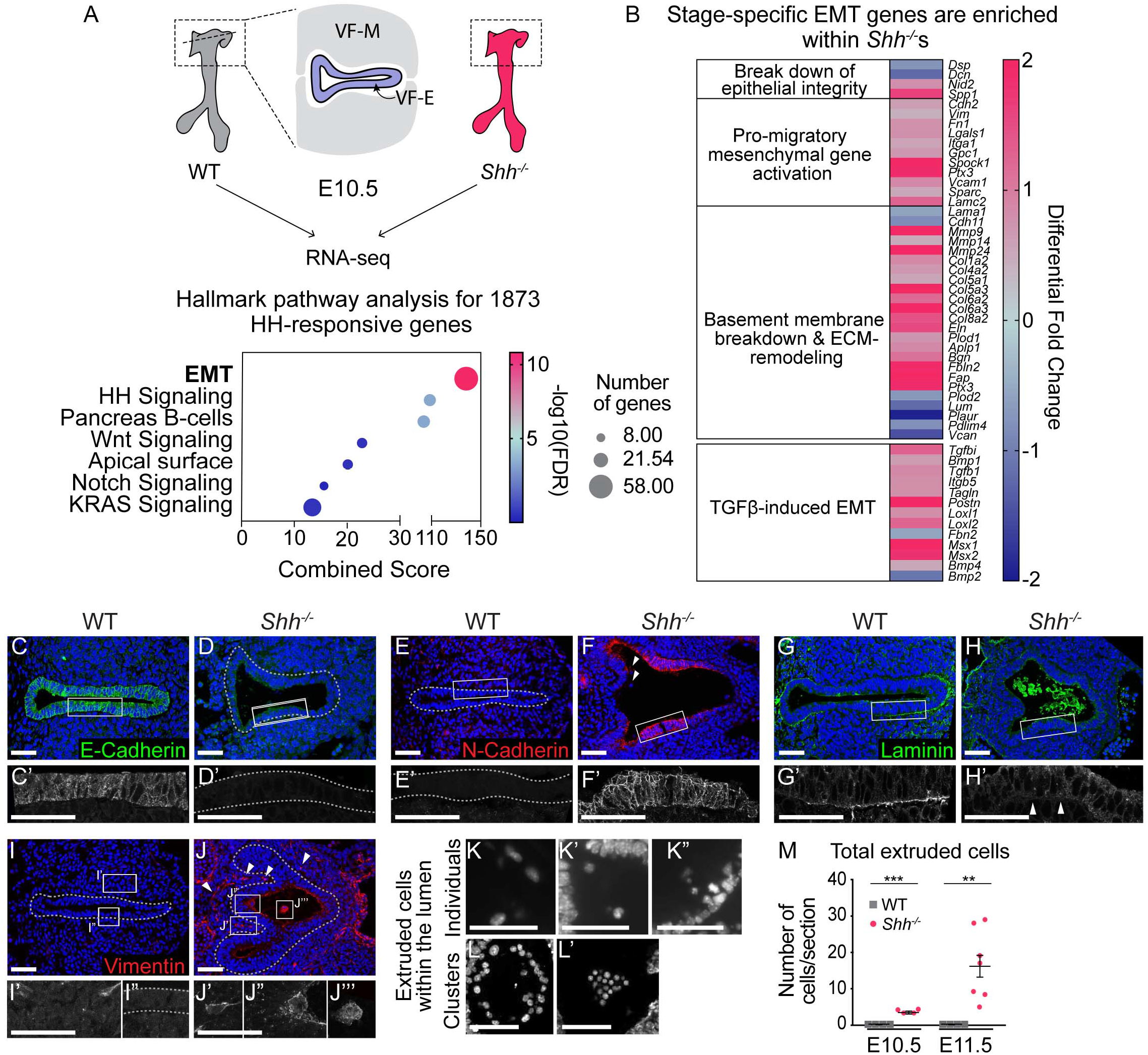
Larynx epithelial cells undergo ectopic EMT-like cell extrusion in the absence of HH signaling. A. RNA-seq of wild-type (WT) and *Shh^-/-^* larynx tissue at E10.5 identified 1873 HH-dependent genes (p<0.05). Epitheli- al-to-mesenchymal transition (EMT)-related genes were highly enriched among HH targets by Hallmark pathway analysis. B. Differentially expressed EMT-genes cluster into stage-specific groups. C-D. E-Cadherin (green) expression in the epithelium across 3 controls and 3 *Shh^7^s* at E10.5. E-F. N-Cadherin (red) expression within the epithelium of 3 controls and 3 *Shh^-/-^s.* Arrowheads mark cells in the lumen. G-H. Laminin (green) expression marking the basement membrane in 3 controls and 3 *Shh+s.* Arrowheads indicate loss of Laminin from the basement membrane in *Shir’s.* I-J. Vimentin (red) expression to mark migrating cells in 3 control and 3 *Shir’-* larynxes at E10.5. Arrowheads mark Vimentin-expressing cells within the lumen (J”, J’”) and the surrounding mesenchyme (J’) in *Shir’s.* K-M. DAPI staining marking cells within the lumen of the larynx in *Shir’s* at E10.5 and E11.5. M. Total number of luminal cells/section were quantified in 4 control and 4 *Shir’s* at E10.5 and in 4 controls and 7 *Shir’s* at E11.5. Average numbers of luminal cells/section were analyzed for statistical significance using the Student’s t-test. Error bars show the standard error of the mean.**p<0.005 ***p<0.0005. VF-M-vocal fold mesenchyme; VF-E-vocal fold epithelium. All scale bars denote 50pm.

Consistent with the RNA-seq data and evocative of the observations in remodeling epithelia (Figure 1), N-Cadherin (*Cdh2*) was highly upregulated within the mutant epithelium along with a substantial loss of pro-epithelial E-Cadherin (*Cdh1*) (Figure 2C-F). This suggested that the adhesive properties of the larynx epithelium were taking on a mesenchymal profile. This change was further accompanied by a loss of Laminin (*Lama1*) from the basement membrane along the epithelium (Figure 2B,G-H) indicating that HH signaling is required to maintain the integrity of this structure. Additional basement membrane component genes such as *Col4a2* and *Nid2* were upregulated by RNA-seq, suggesting that loss of Laminin may result in a compensatory increase in other basement membrane components in order to maintain membrane integrity (Jones et al., 2016; Salmivirta et al., 2002) (Figure 2B). This was further supported by the persistence of a morphologically intact epithelial layer in *Shh^-/-^* embryos at later stages of larynx development (Figure 2-figure supplement 1A-B).

In some systems, the expression of Vimentin is necessary and sufficient to induce EMT (Huang et al., 2012; Liu et al., 2015; Mendez et al., 2010; Vuoriluoto et al., 2011). As suggested by the transcriptional increase in pro-migratory factors, there was an increase in the level of Vimentin expression both within the epithelium as well as in several cells within the surrounding mesenchyme in *Shh^-/-^*s (Figure 2I-J). High levels of Vimentin expression also marked cells that appear to be extruding from the epithelial layer into the lumen (Figure 2J”, J’’’), as well as clusters of extruded cells within the lumen of the epithelium (Figure 2K’, K”). These clusters were first seen at E10.5 and increased dramatically by E11.5 (Figure 2K-M). Overall, these observations are consistent with laryngeal epithelial cells undergoing EMT in the absence of HH signaling.

### HH signaling prevents a cadherin-switch within the epithelium during early stages of foregut development

To determine the onset of this phenotype, we examined earlier stages of foregut development and found a reduction in the levels of E-Cadherin in the *Shh^-/-^* epithelium of the presumptive larynx as early as E9.5 (Figure 3A-B). While N-Cadherin was not expressed at high levels, expression of membranous N-Cadherin was first observed in a small number of cells within the epithelial layer at E9.5 and at E9.75 (Figure 3C-D, H-I, N), suggesting that N-Cadherin was beginning to be expressed at the same time that mutant epithelial cells began to downregulate E-Cadherin. This is consistent with recent studies that have described co-expression of E-Cadherin and N-Cadherin in cells undergoing EMT (Ray et al 2016, Aiello et al 2018). Over the next few hours of development E-Cadherin re-localized within cells, accumulating in puncta along the apical surface (Figure 3E-G), and was almost completely gone by E10.0 (Figure 3J-K) (Aiello et al., 2018; Woichansky et al., 2016). At this stage, more than half of the cells within the epithelium expressed robust levels of membranous N-Cadherin (Figure 3L-N).

**Figure 3.**
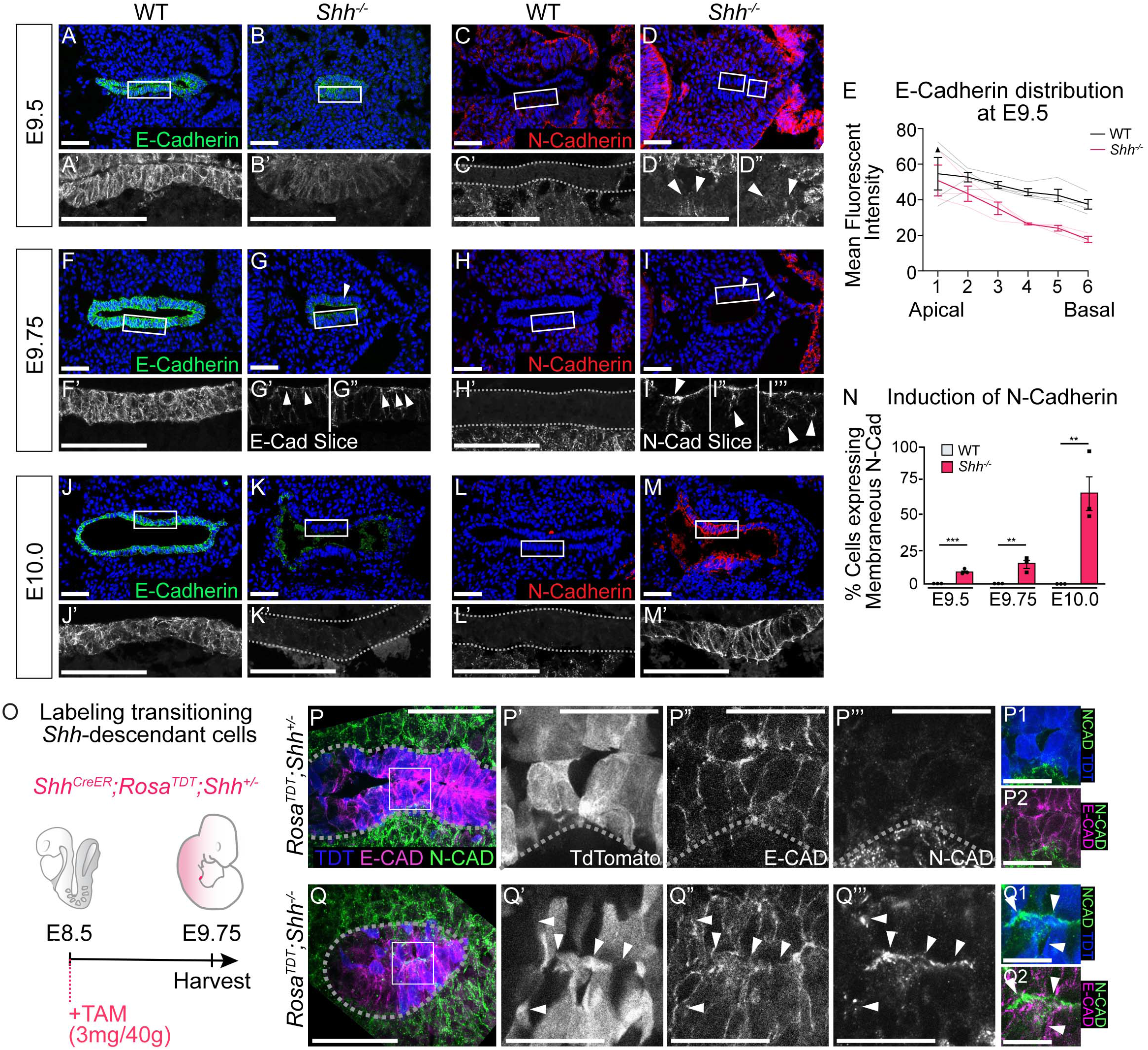
HH signaling is required to prevent a cadherin-switch within the epithelium during early stages of foregut development. A-D. E-cadherin expression (A-B; green) and N-cadherin expression (C-D; red) in 4 control and 3 *Shh^-/-^* larynxes at E9.5 (24-26 somites). E. E-Cadherin distribution along the apical-basal axis of the epithelium in 4 controls and 3 *Shh^7^s* at E9.5. Error bars show standard error of the mean. F-l. E-cadherin expression (F-G; green) and N-cadherin expression (H-l; red) in 3 control and 3 *Shir^7^’* larynxes at E9.75 (27-29 somites). J-M. E-cadherin expression (J-K; green) and N-cadherin expression (L-M; red) was examined in 3 control and 3 *Shh^-/-^* larynxes at E10.0 (29-31 somites). N. The percentage of N-Cadherin-expressing cells within the epithelium at E9.5, E9.75 and E10.0 was averaged across 3 controls and 3 *Shh-^-/-^* larynxes at each stage and analyzed for significance using the Student’s t-test (**p<0.005 ***p<0.0005). Error bars indicate the standard error of the mean. O. Shh-fate mapping in *Shh^CreER^;Rosa^TDT^;Shh^+/^-* and *Shh^CreER^;Rosa^TDT^;Shh’^-/-^* embryos (4 replicates each). Td-Tomato (TDT) labeling was induced with Tamoxifen at E8.5 and visualized at E9.75. P-Q. Sections were analyzed for E-Cadherin (E-CAD; magenta) and N-Cadherin (N-CAD; green) expression as well as Td-Tomato (TDT; blue) expression. Arrowheads mark regions of E-Cadherin and N-Cadherin expression along the membrane of a Td-Tomato-positive cell. P’-P2,Q’-Q2. Scale bars denote 25pm. All other scale bars denote 50pm.

The change in cadherin status suggested a transition to a mesenchymal fate in the absence of HH signaling. Alternatively, these cells might be replaced by a different population of N-Cadherin expressing cells. To distinguish between these possibilities we examined Cadherin expression in larynx epithelial cells using the *Shh^CreER^*;*Rosa^TDT^* reporter line to label *Shh*-expressing epithelial cells in control (*Shh^CreER/+^*) and mutant *Shh^CreER/-^* embryos. Tamoxifen induction at early stages of foregut development (E8.5) exclusively labeled epithelial cells in the vocal folds at E9.75 in control (*Shh^CreER/+^*) and mutant (*Shh^CreER/-^*) embryos (Figure 3P-Q). While E-Cadherin and N-Cadherin had mutually exclusive boundaries restricted to the epithelium and mesenchyme respectively in controls, they appeared to be co-expressed within a small number of TDT-expressing epithelial cells in *Shh^-/-^*s at E9.75 (Figure 3 P-Q; Figure 3-figure supplement 1A-B,D). Co-expression was also observed at E10.0, both as distinct apical puncta as well as laterally along cell-cell boundaries (Figure 3-figure supplement 1C), indicating that some laryngeal epithelial cells undergo a cadherin switch in the absence of HH signaling. The switch in cadherin expression within the vocal folds also occurred on the transcriptional level with *Cdh1* expression in the mutant epithelium at E9.25 replaced by high levels of *Cdh2* expression by E10.5 (Figure 3-figure supplement 2A-D). We conclude that HH signaling is required for the transcriptional maintenance of *Cdh1* along the early foregut.

FOXA2 activates the expression of *Cdh1* and suppresses EMT programs in the endoderm (Bow et al., 2020; Scheibner et al., 2021; Zhang et al., 2015). This suggested that EMT initiation within *Shh^-/-^*s might be caused by a loss of FOXA2 expression. Consistent with this notion, FOXA2 was expressed in nearly every cell at the region of the future larynx in both control and *Shh^-/-^*s during early stages of foregut development (E9.25; 21-23 somites; Figure 4A, Figure 4-figure supplement 1A-B). However, by E9.75 (27-29 somites), FOXA2 expression was absent from ∼30-40% of the mutant cells with reduced expression in many of the remaining FOXA2+ cells (Figure 4A-C, Figure 4-figure supplement 1C-F). FOXA2 was further reduced by E10.5 and completely absent by E11.5 (Figure 4A, Figure4-figure supplement 1G-J). This suggested that HH might prevent EMT by positively regulating FOXA2 in either a cell non-autonomous or autonomous fashion. In keeping with the latter possibility, there was *Ptch1* and low level *Gli1* expression within the larynx epithelium (Figure 4-figure supplement 2A-C), indicating that HH signaling could potentially regulate FOXA2 through autocrine signaling.

**Figure 4.**
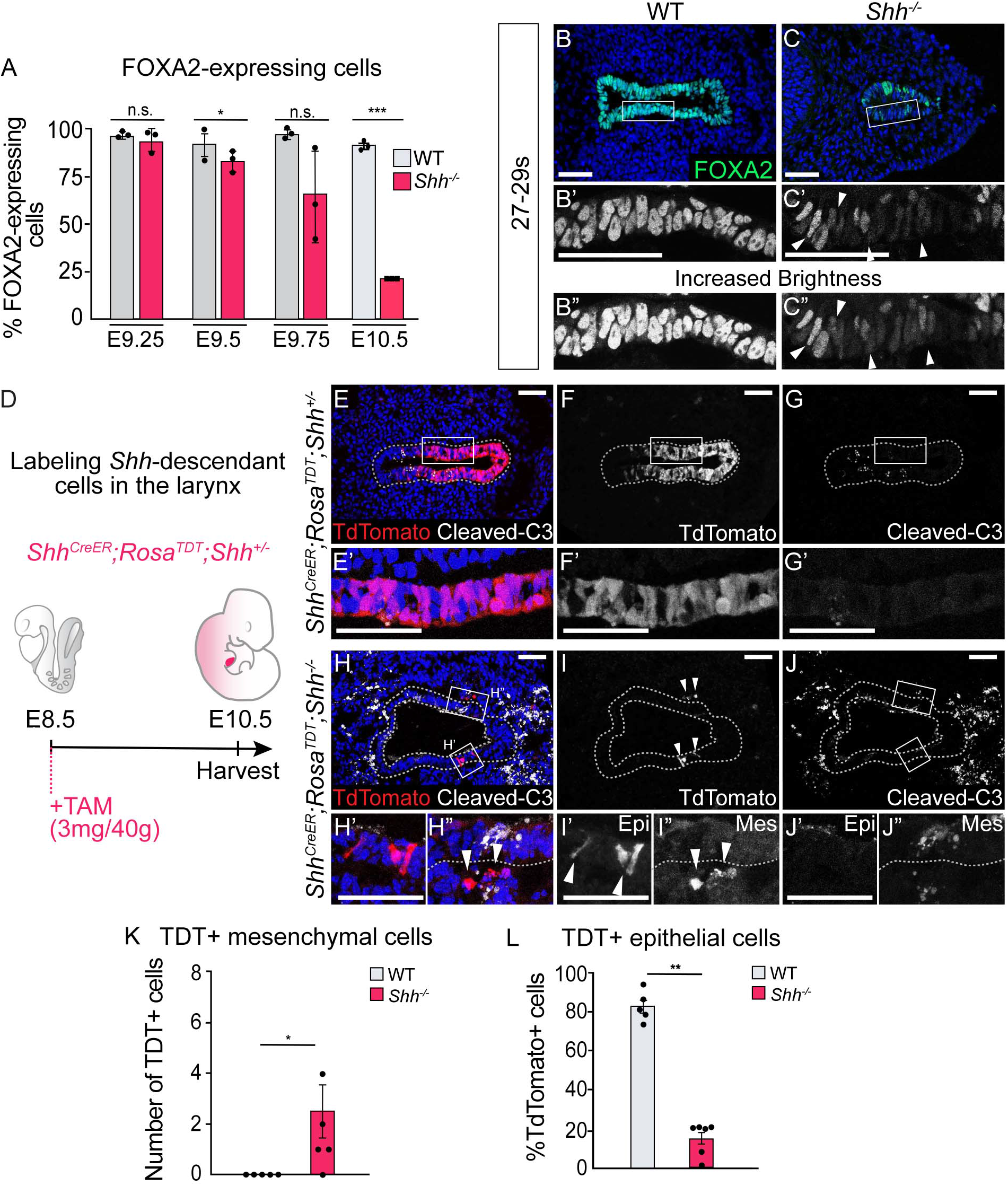
Epithelial cells lose FOXA2 and leave the epithelium in the absence of HH signaling. A. F0XA2 expression in control and *Shh^7^∼* larynxes at E9.25 (21-23s), E9.5 (24-26s), E9.75 (27-29s) and E10.5 (32-35s). The percentage of epithelial cells expressing F0XA2 in 3 control and 3 *Shir^7^’* larynxes was averaged at each develop­mental stage and analyzed for significance using a Student’s t-test. Error bars show the standard error of the mean. B-C. F0XA2 (green) expression is reduced in *Shtr^7^∼s* by E9.75 (27-29s) compared to controls. Arrowheads indicate FOXA2-low cells (B’,B”). Panels B and C have been repeated in Supplemental Figure 4 for clarity. D-J. 5 Shh^CreER^ ;Rosa^TDT^;Shh^-/-^ (E-G) and 5 Shh^CreER^;Rosa^TDT^;Shh^-/-^ (H-J) embryos were induced with Tamoxifen at E8.5 and ana­lyzed for TD-Tomato (TDT) expressing Shh-descendant cells (in red; E-F, H-l) and for Cleaved-Caspase3 expres­sion (in white; E,G,H,J) in the larynx at E10.5. Arrowheads indicate Shh-descendant cells within the epithelium (I’) and within the mesenchyme (H”,l”). K. The number of TDT-expressing cells found in the mesenchyme in 5 controls and 5 *Shh^-/-^* was quantified and tested for significance using the Student’s t-test. Error bars show the standard error of the mean. L. The percentage of TDT-expressing cells within the ventral half of the epithelium in 5 controls and 5 Shh^-/-^s was quantified and tested for significance using the Student’s t-test. Error bars show the standard error of the mean. *p<0.05 **p<0.005 ***p<0.0005; n.s.-not significant. All scale bars denote 50pm.

### Transitioning epithelial cells extrude from the epithelial layer and undergo apoptosis in the absence of HH signaling

We next asked what happened to foregut epithelial cells undergoing EMT once they left the epithelium. These cells could be in the process of undergoing apoptosis, as often happens with extruded cells (Fadul and Rosenblatt, 2018; Kim et al., 2015; Kuipers et al., 2014; Ohsawa et al., 2018). Alternatively, these cells might persist in the mesenchyme adjacent to the epithelium. To address this, we again used genetic fate mapping to examined the fate of larynx epithelial cells, in control (*Shh^CreER/+^*) and mutant *Shh^CreER/-^* embryos (Figure 4D). While Td-Tomato labeling remained restricted to the epithelial layer in control embryos (Figure 4E-G, K) there were a small number of labeled cells within the mesenchyme surrounding the epithelium in mutant embryos by E10.5 (Figure 4H-I, K). These cells, which were relatively rare, were consistent with EMT induction, and the basal extrusion of epithelial cells in the absence of HH signaling (Figure 4H-I, K). Nonetheless, the low number of these cells suggested that most of the cells leaving the epithelium do not survive. Consistent with this idea, there were high levels of cell death in both the mesenchymal and epithelial tissues of the vocal folds between E9.5-11.5, peaking at over 30% of the epithelium (Figure 4H,J; Figure4-figure supplement 3A-H). Moreover, the majority of *Shh^CreER/-^*;*Rosa^TdT^* labeled cells outside the epithelium expressed the apoptosis marker Cleaved Caspase-3 (Figure 4H,J). We conclude that most of the vocal fold cells undergoing EMT in *Shh^-/-^* embryos are either in the process of undergoing apoptosis or undergo apoptosis shortly after extrusion.

### Initial *Shh*-expressing epithelial cells are replaced by an aberrant cell population in the absence of HH signaling

During the initial period of cell death, proliferation levels remained unchanged. However, by E11.5 there was a significant increase in cell proliferation within the vocal fold epithelium of *Shh^-/-^* embryos (Figure 4-figure supplement 3I-J). This, and the persistence of a morphologically distinct epithelium, implied that HH-independent mechanisms might contribute to epithelial maintenance. Notably, the *Shh^-/-^* epithelium was highly disorganized. Compared to the uniform, 1-2 cell layers observed in control embryos, mutant embryos had highly variable epithelia containing increased numbers of cell layers (an average of 12 layers; Figure 4-figure supplement 4A-C), with an overall thickening of the vocal fold epithelium. This aberrant epithelium continued to persist until at least E13.5, and was composed of rudimentary, poorly keratinized, p63-negative cells that do not recover normal epithelial form or function (Figure 2-figure supplement 1A-B).

We asked if the epithelial cells that persist to later stages are descendants of the initial cells marked by *Shh*. Using the same *Shh^CreER/-^*;*Rosa^TdT^* embryos described above, we found that *Shh*- descendants were primarily localized to the ventral half of the vocal fold epithelium by E10.5 (Figure 4D-F, L). As *Shh* is initially expressed throughout the foregut and continues to be expressed within the dorsal larynx epithelium at E11.5, this suggests that foregut epithelial cells undergo dynamic regional cellular rearrangements during this timepoint (Burke and Oliver, 2002; Lungova et al., 2018, 2015; Moore-Scott and Manley, 2005; Motoyama et al., 1998; Rankin et al., 2016; Sagai et al., 2009; Szabo et al., 2009). In marked contrast to control embryos, there was a 60% reduction in *Shh*-descendant cells in *Shh^CreER/-^* mutants (Figure 4H-J, L), indicating that the aberrant epithelium observed at later stages was not descended from the initial epithelium. However, as *Shh*-descendant cells only labeled the ventral half in these experiments, this result did not rule out the possibility that a domain of dorsally located cells repopulated the ventral epithelium. To address this more directly, we examined the expression of SOX2 and NKX2.1, which respectively specify dorsal and ventral foregut domains (Kim et al., 2019; Kuwahara et al., 2020; Nasr et al., 2019; Que et al., 2007). At E10.5 there is an absence of NKX2.1 and a significant reduction in SOX2, which is undetectable by E11.5 (Figure 5A-E, Figure 5- figure supplement 1A- D). We conclude that HH is required for the survival and regional identity of ventral and dorsal endodermal cells that together comprise the epithelium.

**Figure 5.**
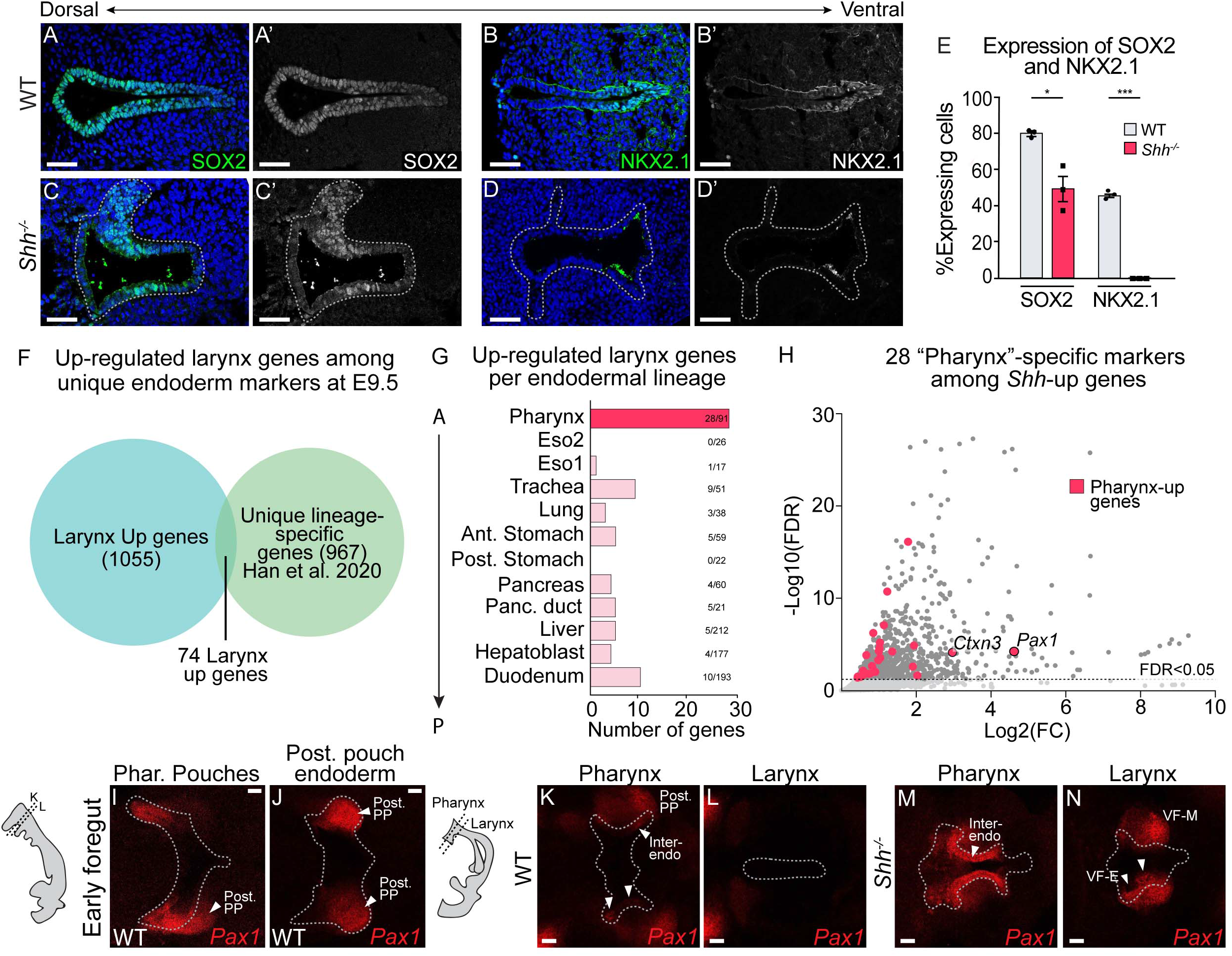
Sbb-descendant larynx epithelial cells are replaced by an ectopic population of Paxf-expressing cells in the absence of HH signaling. A-E. S0X2 (A,C) and NKX2.1 (B-D) expression in 3 control and *Shh^7^’* larynxes at E10.5 (n=3 per genotype). E. The percentage of SOX2 and NKX2.1-expressing cells within the epitheli­um was quantified in 3 controls and 3 *Shir’s* each at E10.5 and analyzed for significance using the Student’s t-test (*p<0.05 ***p<0.0005). Error bars plot the standard error of the mean. F. Endodermal genes identified by Han et al. 2020 were used to infer the composition of the epithelium in *Shir^7^’* embryos. Up-regulated genes within the *Shlr^-/-^* RNA-seq dataset (1056) were intersected with lineage specific endodermal genes (968) identified at E9.5 (Han et al., 2020) resulting in 74 annotated endodermal genes that were up-regulated in the absence of HH signaling. G. Lineage-specific categorization of the 74 up-regulated genes in the larynx showed that pharynx-specific genes (28/74) are enriched within this group. H. The expression levels of the 28 pharynx-specific genes (pink) among all upregulated genes within *Shir^-/-^* larynxes (grey). The dashed line denotes the FDR cutoff <0.05. I-J. *Pax1* (red) expression within the pharyngeal pouches in control E9.5 foregut tissues (n=3). K-N. *Pax1* (red) expression within the pharynx and larynx of control (K-L) and *Shir^-/-^* (M-N) embryos at E10.5 (n=3 for each genotype). Post. PP-poste rior pharyngeal pouch; Inter-endo.-inter-pouch endoderm; VF-M-vocal fold mesenchyme; VF-E-vocal fold epitheli­um. All scale bars denote 50pm. For the list of genes used for this intersection refer to Supplemental Data Table 2.

To determine the identity of the remaining cells that make up the epithelium in the absence of HH signaling, we intersected significantly up-regulated genes found in the E10.5 *Shh^-/-^* larynx RNA- seq dataset (Figure 2A) with a published scRNA-seq dataset of lineage-specific genes generated in the E9.5 endoderm (Figure 5F; Supplemental Data Table 2) (Han et al., 2020). While some lineage specific markers throughout the anterior-posterior axis of the gut were increased in the *Shh^-/-^* dataset (Figure 5G), over a third of the upregulated genes within this intersection (28/74) were annotated as pharyngeal genes. Pharynx-specific genes were significantly more likely to be upregulated in *Shh^-/-^* larynx tissue than trachea/esophagus-specific genes (p=3.215e-05, one-tailed Fisher’s exact test), a finding that was confirmed by gene-set enrichment analysis (Figure 5-figure supplement 2A-B). *Pax1* was the most upregulated pharynx-specific gene within the *Shh^-/-^* dataset (Figure 5H). In controls at this stage, *Pax1* was expressed exclusively in the pharyngeal pouches, which did not express FOXA2 or *Shh* (Figure 5I-J, Figure 5-figure supplement 2C,E,M) (Johansson et al., 2015; Moore-Scott and Manley, 2005; Westerlund, 2013). By E10.5, following larynx specification, *Pax1* continued to be expressed in the *Shh*-negative pharyngeal pouches, and was completely absent from the epithelium and mesenchyme of the vocal folds (Figure 5K-L) (Moore-Scott and Manley, 2005). In contrast, *Shh^-/-^* embryos had ectopic *Pax1* expression throughout the anterior foregut endoderm and larynx. Ectopic *Pax1* expression was also detected in the adjacent mesenchyme of the larynx but was excluded from more posterior foregut tissues such as the trachea and esophagus (Figure 5M-N; Figure 5-figure supplement 2C-L). Correspondingly, FOXA2 expression was either severely reduced or absent from these regions by E11.5 (Figure 5-figure supplement 2M-N). It is presently unclear whether this *Pax1*-expressing population represents an expansion of *Pax1*-expressing cells from the pharyngeal pouches, an expansion of a rare population of cells that is not detectable in controls, or an abnormal population that is not normally found in embryos. We conclude that the initial foregut epithelium is replaced by a different population of *Pax1*-expressing cells. Notably, this indicates a requirement for *Shh* in maintaining epithelial identity that extends beyond the larynx to other parts of the anterior foregut.

## Discussion

We have uncovered a broad role for HH signaling in regulating the morphogenesis of the laryngeal epithelium and propose that this extends to the morphogenesis of additional anterior-foregut-derived organs, including trachea-esophageal separation. There is an unexpectedly early role for HH signaling in maintaining the nascent foregut epithelium, which in its absence undergoes EMT marked by cadherin switching, cell extrusion and ultimately cell death. As this initial population of epithelium dies, it is replaced by an ectopic population of cells (Figure 6A). The unexpected presence of this population complicates the previous interpretation of HH mutant phenotypes in the anterior foregut, as changes in gene expression that have been interpreted as reflecting HH-dependent transcriptional changes might instead reflect the properties of the new population of replacement cells.

**Figure 6.**
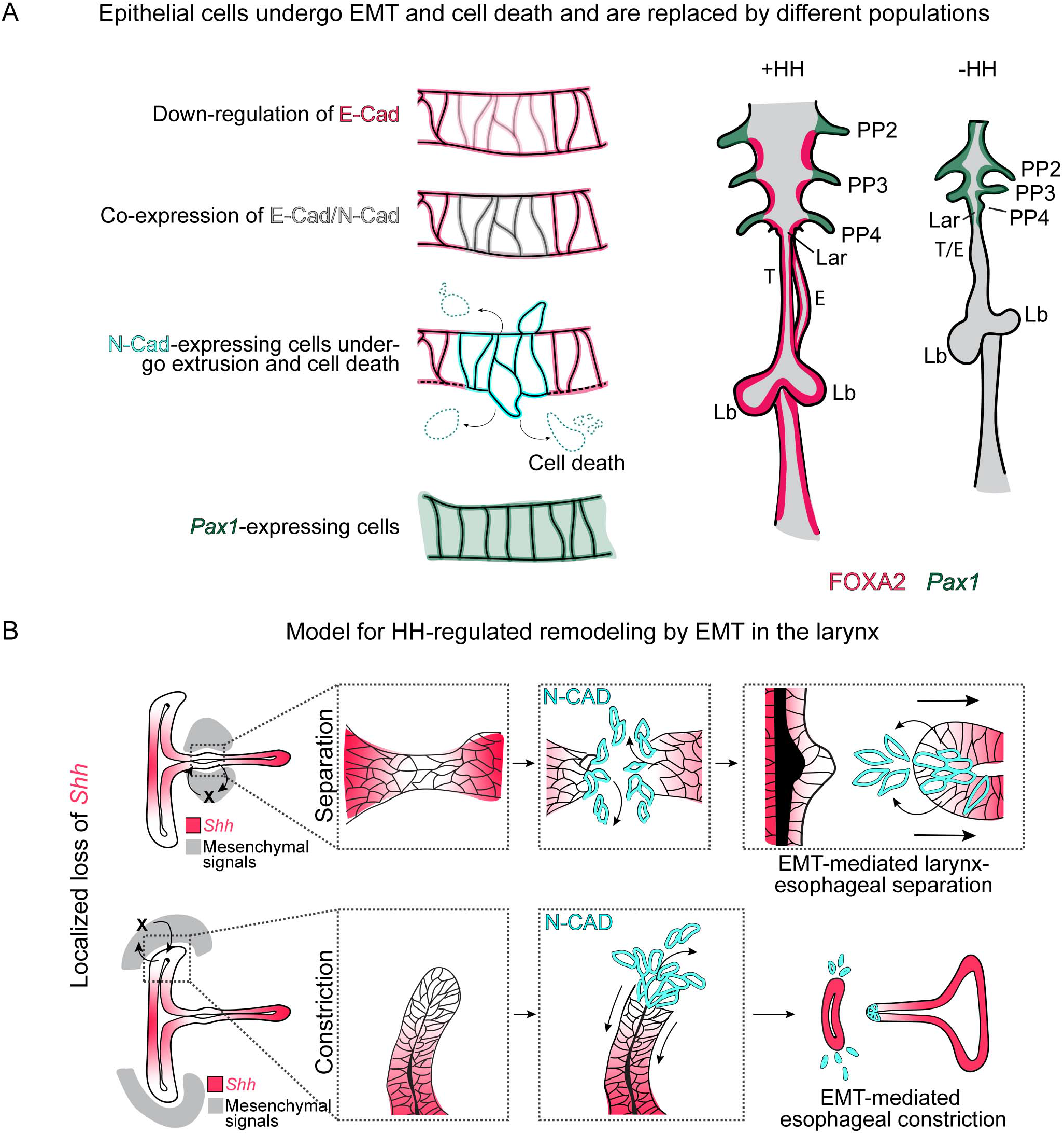
Dynamic HH signaling drives the homeostasis of the early anterior foregut endoderm and may also regulate later stages of larynx remodeling. A. Epithelial cells marked by E-Cadherin and FOXA2 (pink) in the anterior foregut undergo EMT (marked by N-Cadherin expression in cyan), migrate out of the foregut epithelium, and undergo cell death in the absence of HH signaling. These cells are then replaced by different Paxf-expressing (dark green) populations. B. We hypothesize that dynamic SHH (pink) expression within the larynx epithelium drives EMT-mediated morphogenesis are later stages of larynx development, specifically larynx-esophageal separation and esophageal constriction. PP 2/3/4-Pharyngeal pouches #2-4, Lar-larynx, T-trachea, E-esophagus, Lb-lung buds.

### The anterior foregut epithelium consists of a new population of cells with aberrant gene expression

The loss of FOXA2, SOX2 and NKX2.1 (Figures 4,5, Figure 5-figure supplement 1,2) could be due to either the transcriptional downregulation of HH target genes or it could reflect the absence of cells that express these foregut markers. The nearly complete loss of *Shh*-descendant labeling coupled with high levels of cell death (Figure 4; Figure 4-figure supplement 3) is consistent with the latter scenario, suggesting that the descendants of *Shh*-expressing endodermal cells are replaced by a PAX1^+^;FOXA2^-^ population. As *Pax1* normally exclusively marks the pharyngeal pouches, and both *Shh* and FOXA2 are excluded from these regions, these cells are most likely to represent the expansion or ectopic expression of this lineage (Figure 6A; Figure 5-figure supplement 2C,E,M) (Johansson et al., 2015; Moore-Scott and Manley, 2005; Müller et al., 1996; Okada et al., 2016; Wallin et al., 1996; Westerlund, 2013). Elevated levels of *Pax1* have previously been identified in the pharyngeal pouches of *Shh^-/-^* embryos and our findings indicate that this extends through the larynx (Moore-Scott and Manley, 2005; Wallin et al., 1996). Consistent with this tissue representing a pharyngeal pouch population, the loss of *Shh* has been reported to result in ectopic pharyngeal pouch-derived thyroid structures at later developmental stages (Fagman et al., 2004; Westerlund, 2013). We do not however detect the thyroid marker NKX2.1 within the *Shh^-/-^* larynx and it is presently unclear whether this might be due to a delayed onset of the thyroid fate, if the expanded *Pax1*-positive regions contain other pharyngeal pouch-derived cells or if this represents a different population of cells altogether.

### Mechanisms underlying HH-mediated epithelial changes

Prior to being lost from the foregut, FOXA2 expression is reduced in *Shh^-/-^* foreguts as well as the pharyngeal endoderm (Figure 4C, Figure 4-figure supplement 2D,F,H) (Yamagishi et al., 2003). *FoxA2* is a direct transcriptional target of SHH in the neural tube (Oosterveen et al., 2012; Peterson et al., 2012; Sasaki and Hogan, 1994) and the HH-target genes *Ptch1* and *Gli1* are expressed within the foregut epithelium (Figure 4-figure supplement 2A,C). This is consistent with a possibility that HH signaling could have a role in the transcriptional maintenance of *FoxA2* or perhaps other epithelial genes through autocrine signaling (Motoyama et al., 1998; Yamagishi et al., 2003). However, as has been previously noted in other foregut tissues, the expression of HH- responsive target genes is much lower in the epithelium than in the mesenchyme (Figure 4-figure supplement 2C) (Han et al., 2017; Moore-Scott and Manley, 2005; Motoyama et al., 1998; Rankin et al., 2016). Thus, HH signaling could alternatively occur through a more conventional paracrine signaling mechanism to regulate the surrounding mesenchyme, which would in turn regulate the epithelium as previously suggested (Han et al., 2020, 2017; Nasr et al., 2021; Rankin et al., 2016; Yang et al., 2021). Consistent with this possibility, there is widespread cell death in the *Shh^-/-^* mesenchyme that precedes that in the epithelium (Figure 4, Figure 4-figure supplement 3G,H). This results in dramatic changes to the composition of the mesenchyme, including the upregulation of multiple TGFß family members that have well-established roles in inducing EMT as well as antagonizing HH signaling during thymic/parathyroid and pancreas induction (Hebrok et al., 1998; Katsuno et al., 2013; Kim and Hebrok, 2001; Mercado-Pimentel and Runyan, 2007; Moore-Scott and Manley, 2005; Nawshad et al., 2004; Schnaper et al., 2003; Thiery et al., 2009).

FOXA2 has recently been reported to prevent EMT and to regulate the expression of N-Cadherin in the nascent endoderm during gastrulation (Scheibner et al., 2021). SHH-mediated regulation of FOXA2 might represent an extension of this process to the anterior foregut. In contrast to gastrulation, the expression of N-Cadherin coincides with a reduction and re-localization of E-Cadherin in the *Shh^-/-^* anterior foregut (Figures 2, 3). In different contexts, E-Cadherin has been reported to be a direct target of FOXA2 in oral and breast cancer cells (Bow et al., 2020; Zhang et al., 2015). Thus, the reduction of FOXA2 from the endoderm could act to initiate EMT and cadherin switching prior to the onset of cell death. This is consistent with a model where HH prevents precocious EMT in the larynx epithelium by maintaining FOXA2 levels by an unknown mechanism. The subsequent, widespread cell death both within the epithelium and the mesenchyme of *Shh^-/-^* laryngeal tissue likely accounts for the expulsion of *Shh*-descendant epithelial cells from the foregut endoderm, a process that is comparable to cell death-induced cell extrusion in other systems (Kuipers et al., 2014; Ohsawa et al., 2018; Rosenblatt et al., 2001). Curiously the gradual loss of E-Cadherin protein within *Shh^-/-^*s was not observed during larynx- esophageal separation, perhaps indicating differences in HH requirements in the early versus late remodeling larynx (Figure 1; Figure 1-figure supplement 1). Alternatively, this could be due to the dynamic expression of *Shh* at late stages of larynx remodeling and a resulting shorter temporal window for EMT onset during larynx-esophageal separation.

Confirming previous studies, we find that *Shh* is dynamically expressed during larynx development (Lungova et al., 2018, 2015; Sagai et al., 2009). Those areas where *Shh* expression is most reduced correspond to the location of unique N-Cadherin positive domains evocative of the earlier global induction of N-Cadherin in *Shh^-/-^* foregut epithelium (Figures 1,2). In addition to its absence from regions of the larynx epithelium that express N-Cadherin, the relative levels of *Shh* within *Shh*-expressing domains of the epithelium are highly dynamic at later stages, where there is an overall reduction in *Shh* within the dorsal half of the larynx epithelium, which is contiguous with the esophagus, compared to the ventral half (Figure 1B,H). Additionally, lower levels of *Shh* have been reported in the trachea compared to the esophageal epithelium at later stages (Nasr et al., 2021). *Shh* expression in the larynx is regulated by three distinct enhancers that occupy largely non-overlapping regions of activity along the dorsal-ventral axis of the larynx (Sagai et al., 2017, 2009; Tsukiji et al., 2014). While it remains unclear how they are regulated, differential enhancer utilization is a plausible mechanism for regional regulation of *Shh* along the foregut.

### A global role for Hedgehog signaling in anterior foregut organogenesis

We propose that regionalized loss of *Shh* within the anterior foregut triggers partial EMT as a key step in driving the morphogenesis of foregut-derived organs. Alternatively, there may be additional regional factors that are required to activate partial EMT upon withdrawal of HH signaling. HH is locally restricted along the foregut endoderm at the initiation sites of multiple foregut-derived organs including the thymus, the pancreas, the thyroid, and the liver (Apelqvist et al., 1997; Bain et al., 2016; Bort et al., 2006; Fagman et al., 2004; Gordon and Manley, 2011; Hebrok, 2000; Hebrok et al., 1998; Moore-Scott and Manley, 2005; Westerlund, 2013). It is unclear why HH restriction is required in these different contexts and if they share a common mechanism. The liver bud is generated from foregut tissue that lacks *Shh* expression and subsequently undergoes EMT into the adjacent mesenchyme (Bort et al., 2006; Mu et al., 2020). Additionally, loss of *Shh* and the expression of N-Cadherin within the foregut epithelium mark the site of the presumptive dorsal and ventral pancreatic buds, though N-Cadherin is dispensable for the initial stages of pancreatic budding (Esni et al., 2001; Johansson et al., 2010). While the role of *Shh* has not been directly studied in this process, *Hhex* mutants, which fail to undergo EMT of the liver bud also mis- express *Shh* in the epithelium (Bort et al., 2006). Given the role for FOXA2 in regulating EMT in gastrulating endoderm, HH signaling could act either directly or indirectly to maintain FOXA2 (Scheibner et al., 2021). This could include the maintenance of FOXA2 expression/activity or by co-regulating a set of common downstream targets.

## Methods and Materials

### Embryonic Manipulations

All experiments involving mice were approved by the Institutional Animal Care and Use Committee at the University of Texas at Austin (protocol AUP-2019-00233). The *Shh^tm1amc^* null allele (referred to as *Shh^+/-^*) was maintained on a Swiss Webster background (Lewis et al., 2001). The *Cg-Shh^tm1(EGFP/cre)Cjt^* (*Shh^GFP-Cre^*) (Harfe et al., 2004), the *Ptch1^tm1Mps^/J* (*Ptch^LacZ^*) (Goodrich et al., 1997), the *Gli1^tm2Alj^/J* (*Gli1^LacZ^*) (Bai et al., 2002), and the *Shh^CreER/+^;Rosa^TDT/+^* lines (Harfe et al., 2004; Srinivas et al., 2001) were maintained on mixed genetic backgrounds. To label *Shh*- descendant cells, pregnant mice containing *Shh^CreER^;Rosa^TDT^* embryos were injected intraperitoneally with 3mg of Tamoxifen (Sigma Aldrich, T5648-1G) per 40g.

### Gene Expression

RNA was extracted using Trizol reagent (Life Technologies, 10296010) and DNAse-treated. For bulk RNA-seq, vocal fold tissue was dissected from two sets of 3-pooled control and *Shh^-/-^* embryos at E10.5 (32-35s). Libraries were generated using the NEBNext Ultra II Directional RNA library prep kit and single-end sequenced on the Illumina NextSeq 500 platform for ∼40,000,000 reads/sample. Sequenced reads were aligned to the mm10 genome using HISAT2 v2.1.0 and assembled into genes using StringTie v1.3.6 (Pertea et al., 2016, 2015). The data from this RNA-seq is accessible from GEO (accession number GSE190281) and differentially expressed genes are listed in Supplemental Data Table 1.

### Immunofluorescence

All immunofluorescence replicates (denoted by n) refer to independent biological replicates from different embryos. For paraffin embedding, embryos were fixed overnight in 10% formalin, sectioned to 5μm and incubated in three 5 minute washes of boiling 10mM sodium citrate buffer, pH6.0 prior to antibody incubation. For OCT embedding, embryos were fixed for 1hr in 4% paraformaldehyde at room temperature, sucrose protected, embedded in OCT and sectioned to 10μm. Samples were then permeabilized in 0.06% PBST (Triton-X) prior to blocking . Paraffin and OCT sections were blocked in 3% bovine serum albumin (BSA) and 5% normal goat serum/PBST (0.1% Tween-20) for 1hr at room-temperature and, following an overnight incubation in primary antibody at 4°C (see Supplemental Data Table 3 for a list of all antibodies), incubated in secondary antibodies for 1hr at room-temperature. Apoptosis was detected on OCT embedded sections by TUNEL staining, using the In Situ Cell death detection Kit (Roche, 12156792910). All samples were counterstained in 4’,6-diamidino-2-phenylindole (DAPI; Invitrogen, D1306) for 10 minutes at room-temperature and mounted in ProLong Gold Antifade (Thermo fisher Scientific, P36930) prior to imaging. The E-Cadherin-488, N-Cadherin, TDT triple stains (Figure 2) were imaged on a Nikon Eclipse Ti-2 microscope equipped with a 60x, 1.40NA objective; a Visitech iSIM super-resolution confocal scan head; and a Photometrics Kinetix22 camera. All other images were obtained using a Zeiss LSM 710/Elyra S.1 confocal microscope and 10x, 20x, or 63x objectives.

To visualize E-cadherin and N-Cadherin co-expressing cells within the larynx epithelium, OCT-embedded sections were permeabilized, blocked, and incubated in unconjugated N-Cadherin/ goat anti-rabbit Alexa 647 as specified above. Sections were then blocked in Rabbit IgG isotype control (Cell Signaling Technologies, 3900S) (in 5% normal goat serum, 1% Triton-X, PBS) for 1hr at room-temperature. Following an overnight incubation in E-Cadherin-488 at 4°C, samples were washed in 1XPBS, counterstained with DAPI as described above, and mounted in ProLong Gold Antifade. For whole-mount immunofluorescent staining, embryos were processed as described by Nasr et al 2019. To image, wholemount stained embryos were embedded in 1.5% low-melt agarose (Sigma, A5030) cooled to room-temperature, and cleared overnight using Ce3D++ which was prepared with a high concentration of iohexol as described by Anderson et al., 2020.

E-Cadherin localization along the apical-basal axis of the epithelium was measured in Fiji using the average fluorescent intensity of E-Cadherin (normalized to background) within a selected region along the lateral wall of the vocal folds, divided into 6 equal regions from the apical to the basal end of the epithelium. Relative levels of RAB-11, GFP, *Shh* and *Cdh1* along the larynx epithelium was measured in Fiji using a 25- or 35pt-thick line scan that was normalized to background fluorescence.

### Wholemount Fluorescent In situ Hybridization (HCR)

All whole mount fluorescent in situ hybridization replicates (denoted by n) refer to independent biological replicates from different embryos. Wholemount HCR was carried out on whole embryos or cultured larynx explants as previously described in Choi et al., 2018, and in Anderson et al., 2020. Briefly, samples were incubated in 16nM probe overnight at 37°C, and then in 30pmol hairpin per 0.5mL of amplification buffer (Molecular Instruments) overnight at room-temperature. After incubation with the hairpins, samples were washed and counterstained in DAPI overnight as specified by Anderson et al., 2020. Samples were then embedded in low-melt agarose and cleared in CeD3++ as described (Anderson et al., 2020) before imaging. See Supplemental Data Table 3 for list of HCR probes used in the study.

## Acknowledgements

We thank John Wallingford and Dan Dickinson for comments on the paper. We thank Dan Dickinson and Naomi Stolpner, for use of the Nikon Eclipse microscope. We thank Susan Mackem for providing the *Shh^CreER^* line. This work was supported by NIH R01 HD090163 (to SAV and HJ), NIH R01 HD093363 (to AMZ), a Continuing Graduate Fellowship and Provost’s Graduate Excellence Fellowship (to JR), and a TIDES Summer Fellowship (to AEB).

---

**Figure 1-figure supplement 1.**
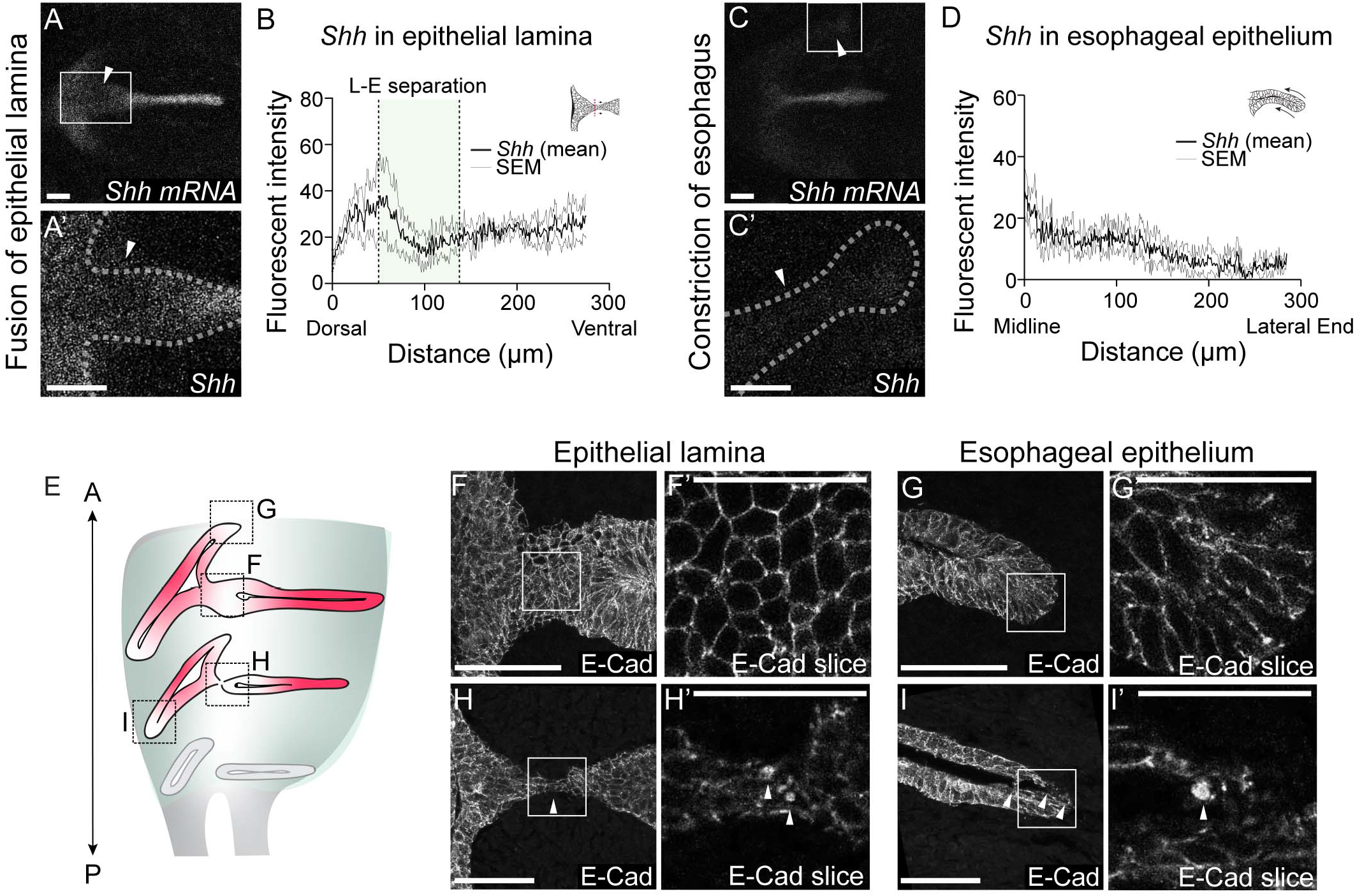
*Shh* expression is reduced, and E-Cadherin is re-localized in the epithelial lamina and the constricting esophagus during larynx-esophageal separation. A-D. *Shh* expression in the epithelial lamina and the constricting esophageal opening in 3 control E11.75 whole-mount larynxes by fluorescent in situ-hybridization. Relative *Shh* (B, D) expression was measured by line scans of fluorescent intensity along the epithelial lamina (A-B) and the esophagus (C-D) in all 3 larynxes. Graphs show average fluorescent intensity and the standard error of mean across the three replicates. E. E-Cadherin protein expression in the epithelial lamina and the constricting esophagus in anterior larynx sections prior to larynx-esophageal separation (F-G), and more posteriorly at the level of larynx-esophageal separation (H-l). E-Cadherin expression and distribution across the cell surface was analyzed in 3 larynxes. H’ and I’. Punctate E-Cadherin expression was observed in single slices at both regions of remodeling at the level of larynx-esophageal separation. A-anterior; P-posterior. F’,G’,H’,r. Scale bars denote 25pm. All other scale bars denote 50pm.

**Figure 1-figure supplement 2.**
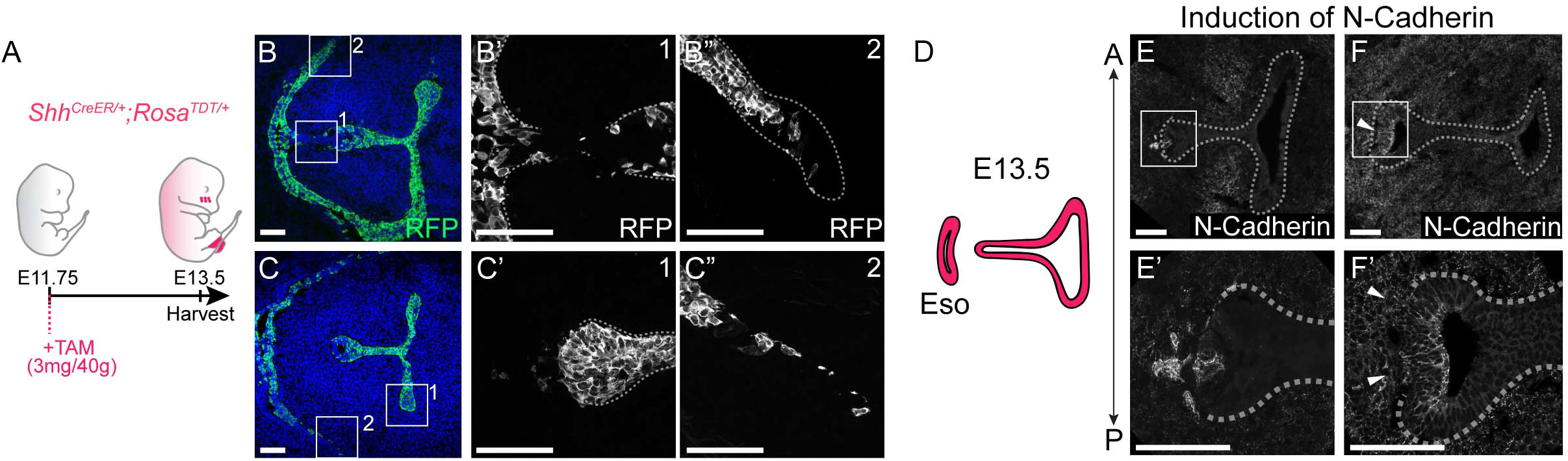
Shh-descendant cells express N-Cadherin and undergo EMT during larynx-esophageal separation. A-C. *Shh^CreER/+^;Rosa^TDT/+^* embryos (3 replicates) were induced with Tamoxifen at E11.75 and examined for RFP (green) expression at E13.5 in anterior (B) and posterior (C) sections of the larynx. Arrowheads indicate extruding RFP-positive Sfrh-descendant cells in the mesenchyme. D-F. N-Cadherin expression in anterior (E) and posterior (F) sections through the larynx at E13.5 at the region of larynx-esophageal separation (3 replicates at each plane). All scale bars denote 50pm.

**Figure 2-figure supplement 1.**
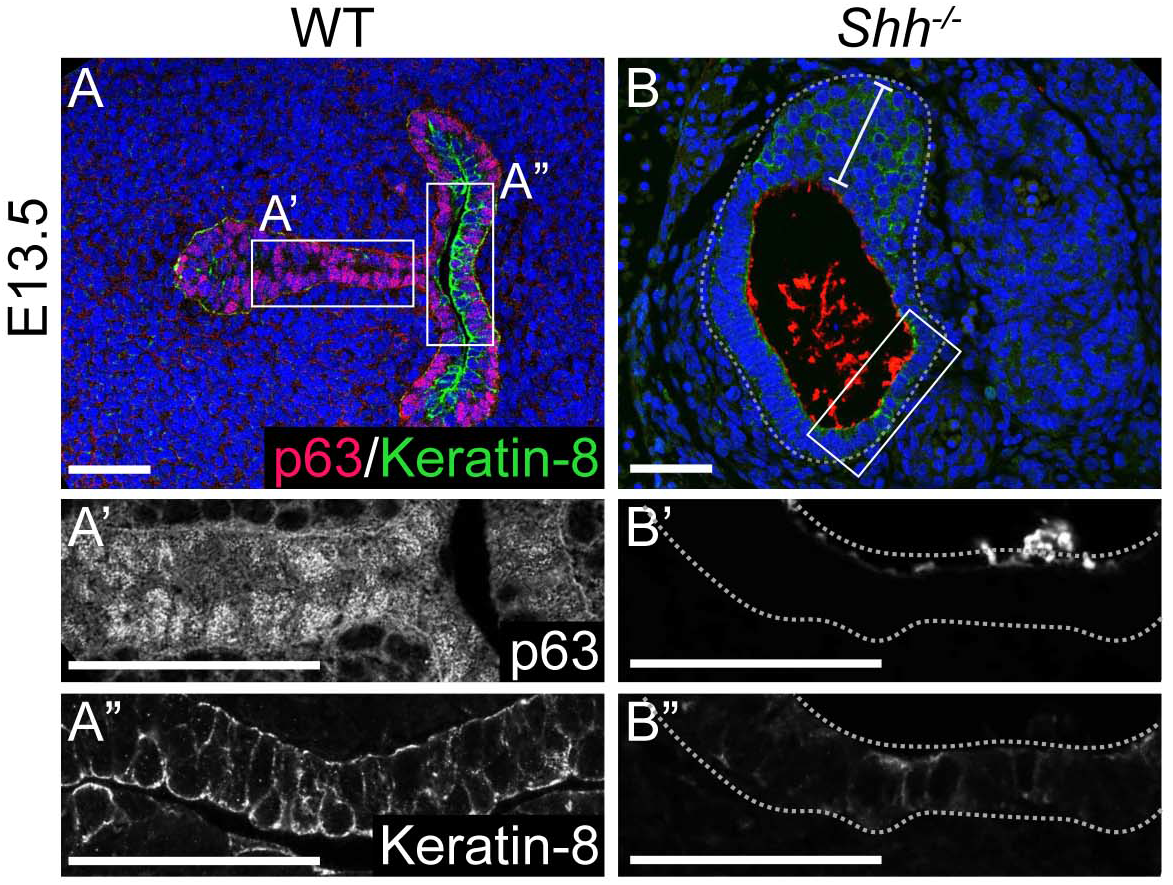
Loss of HH signaling results in the persistence of a poorly keratinized, p63-negative epithelium at late stages of larynx development. A-B. P-63 (red) and Keratin-8 (green) expression in 3 control and 3 *Shh^-/-^* larynxes at E13.5. All scale bars denote 50pm.

**Figure 3-figure supplement 1.**
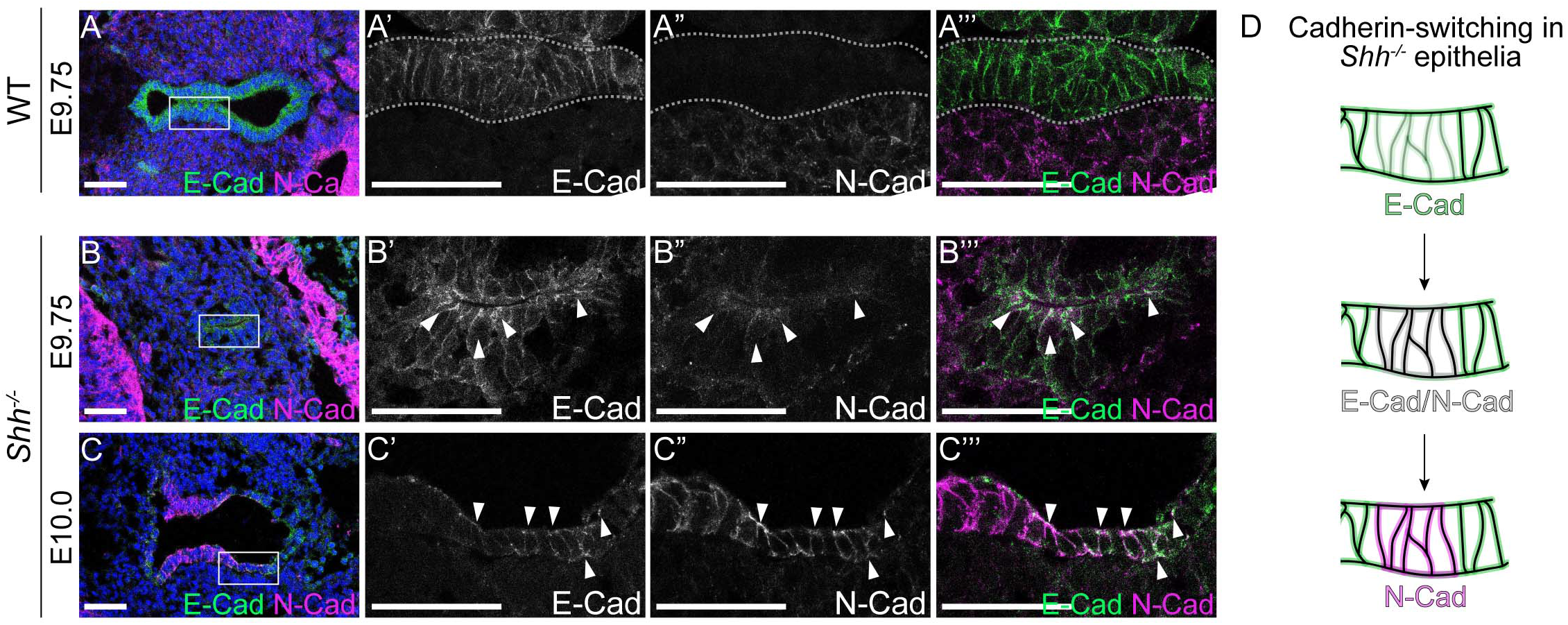
Larynx epithelial cells co-express E-Cadherin and N-Cadherin during early stages of foregut development in the absence of HH signaling. A-D. E-Cadherin (green) and N-Cadherin (magenta) in the vocal fold epithelium of 3 control and 3 *Shh^-/-^* embryos at E9.75 (27-28s) (A-B) and at E10.0 (29-31 s) (C). White arrowheads mark regions of co-expression along the apical surface as well as along the cell boundaries. All scale bars denote 50pm.

**Figure 3-figure supplement 2.**
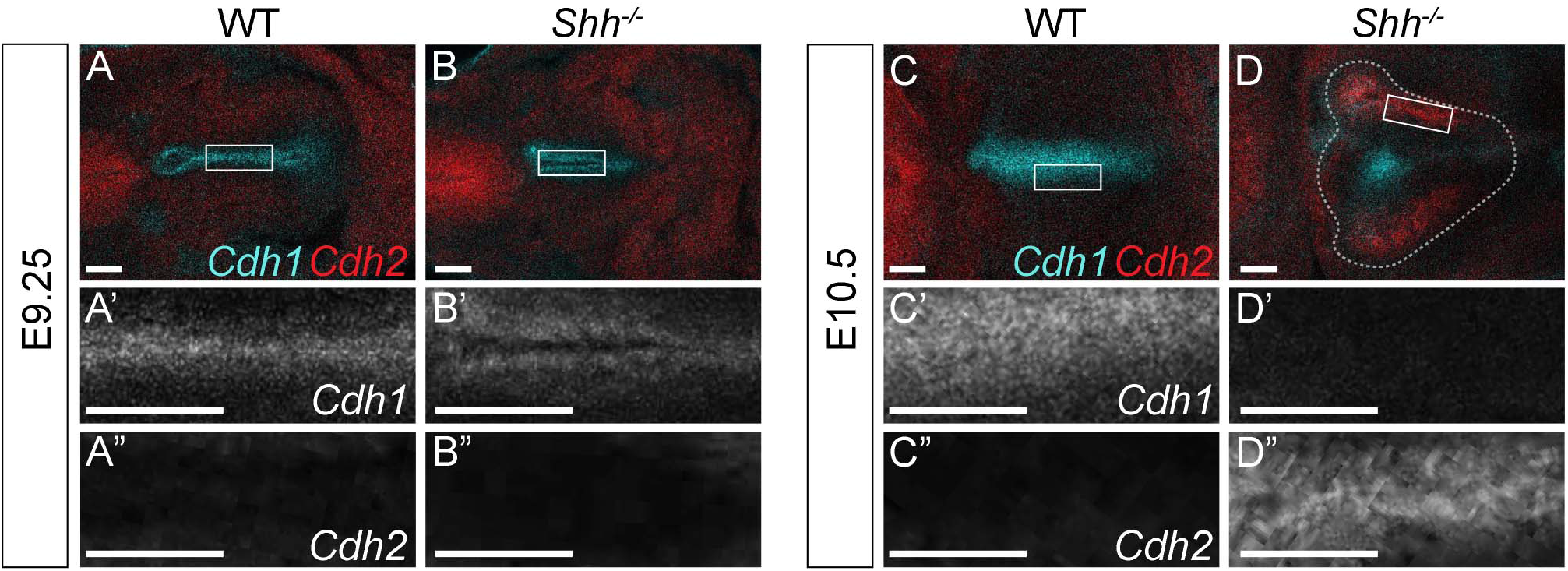
HH is required to maintain *Cdh1* expression in the early foregut. A-D. *Cdh1* (cyan) and *Cdh2* (red) expression by whole-mount fluorescent in-situ hybridization in 3 control and 3 *Shh ^7^’* larynxes at E9.25 (A-B) and at E10.5 (C-D). All scale bars denote 50pm.

**Figure 4-figure supplement 1.**
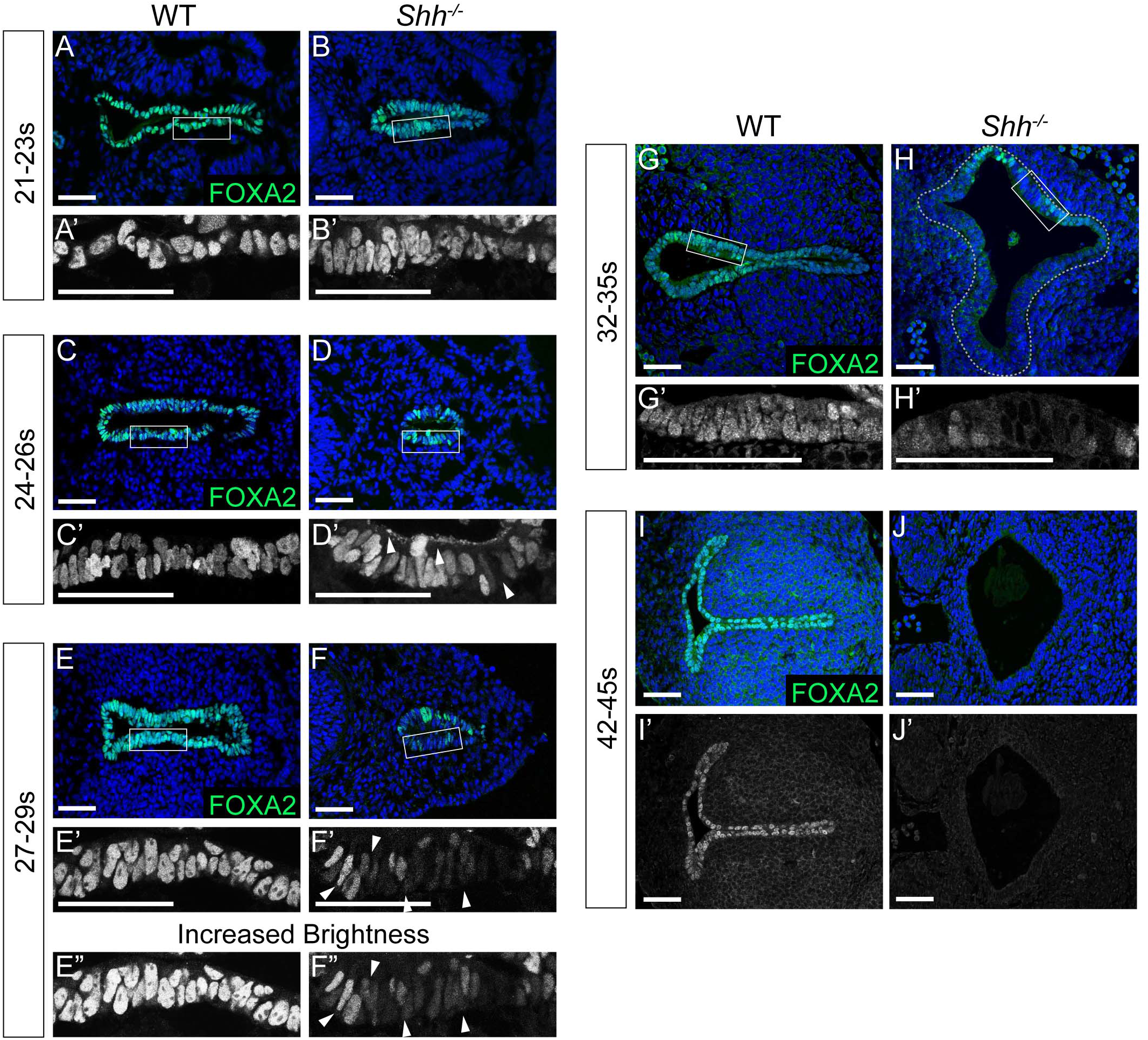
F0XA2 is downregulated in the early foregut endoderm and absent from the larynx epithelium by E11.5 in *Shir’s.* A-J. FOXA2 expression (green) in control and *Shir^7^’* larynxes at E9.25 (21-23S, A-B), E9.5 (24-26s, C-D), E9.75 (27-29s, E-F), E10.5 (32-35s, G-H), and E11.5 (42-45s, l-J). 3 controls and 3 mutants were examined at each timepoint. Panels E and F have been repeated here from Figure 3 for clarity. All scale bars denote 50pm.

**Figure 4-figure supplement 2.**
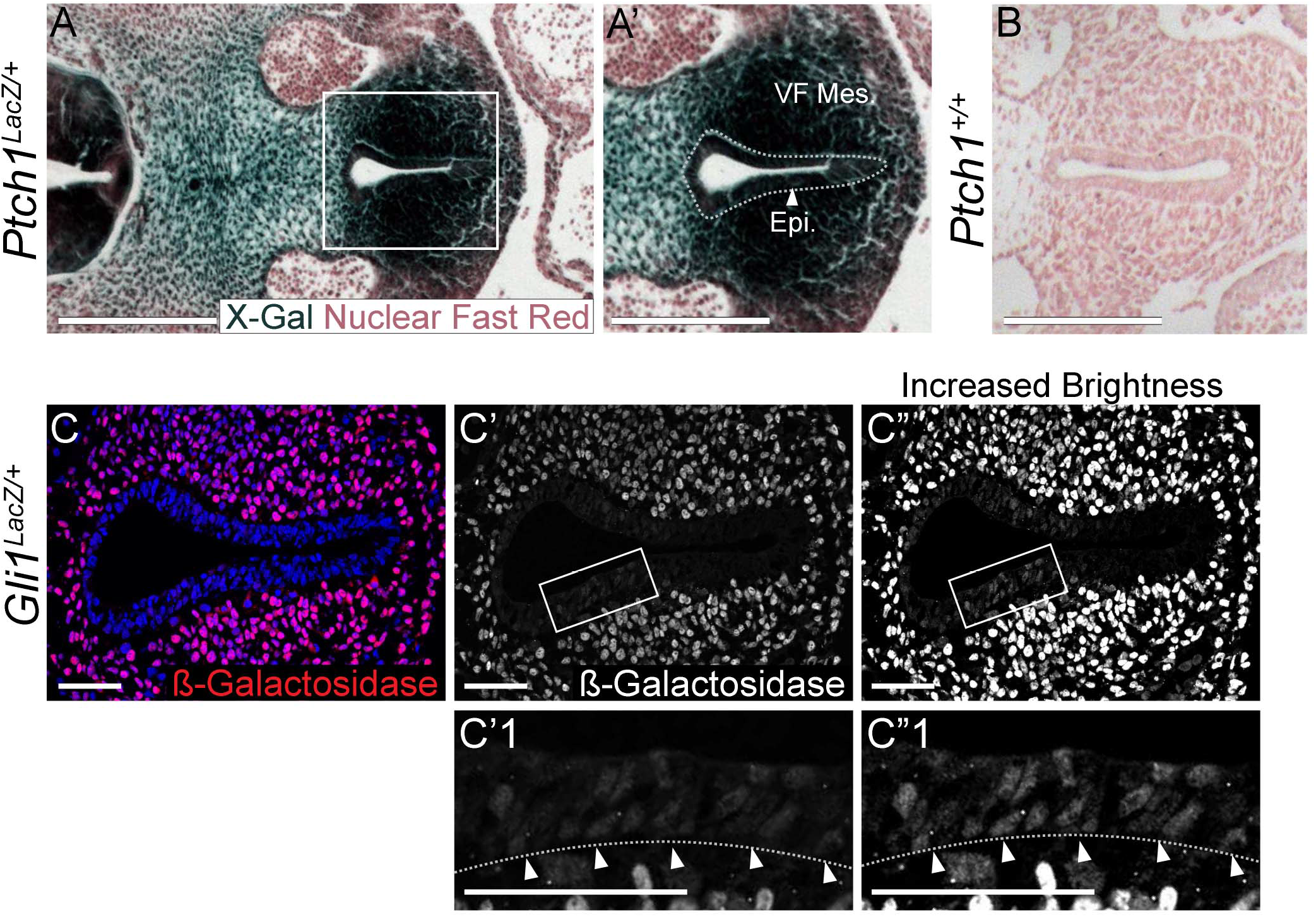
*Ptch^LacZ^* and *Gli1^LacZ^* expression is present within the larynx epithelium as well as the mesenchyme at E10.5. A-A’. *Ptch^LacZ^* expression by ft-galactosidase (marked by X-Gal in dark blue) staining of 3 *Ptch^LacZ/+^* embryos at E10.5. Embryos were fixed in 1% formaldehyde/0.2% glutaraldehyde, stained in 1mg/ml X-gal for 4 hours and sectioned to generate 5 pm sections that were then counterstained with Nuclear Fast Red. B. Sections through the larynx from control *(Ptch^+/+^)* embryos stained for IJ-galactosidase and Nuclear Fast Red. A-B. Scale bars denote 250pm. C-C”. *GH1^LacZ^* expression within the larynx epithelium and mesenchyme of 3 *Gli1^LacZ/+^* E10.0 embryos was assessed using a U-galactosidase antibody (red; Abeam ab9361). Scale bars denote 50pm.

**Figure 4-figure supplement 3.**
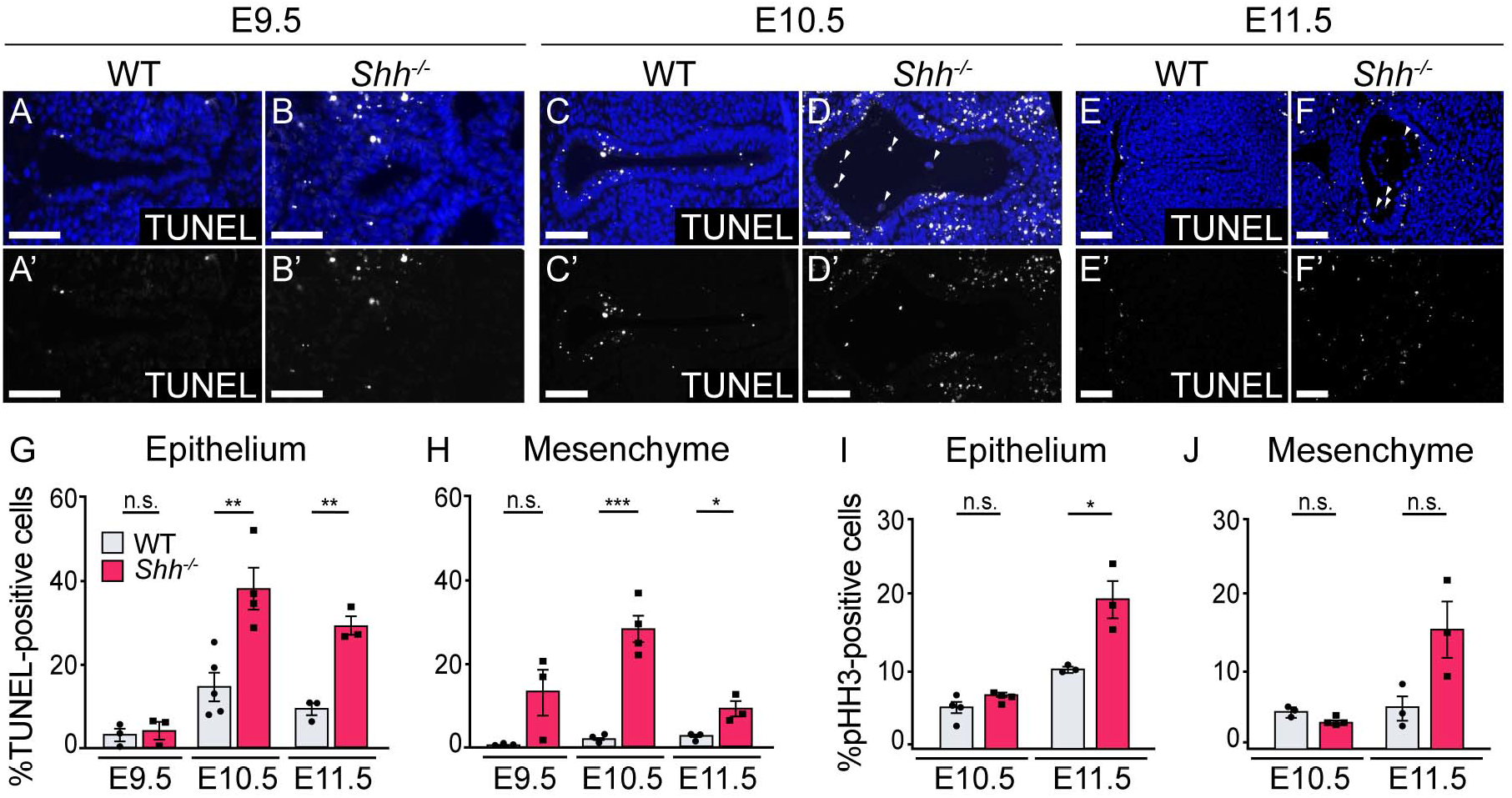
HH signaling is required for the survival of epithelial and mesenchymal cells within the larynx. A-F. TUNEL-staining of control and *Shh^-/-^s* at E9.5 (A-B), 10.5 (C-D) and E11.5 (E-F) within the epithelium and mesenchyme of the vocal folds. G-H. The percentage of TUNEL-positive cells within the epithelium and mesenchyme was quantified over 3-5 replicates each at E9.5, 10.5 and 11.5. I-J. The percentage of proliferative pHH3-positive cells within the epithelium and mesenchyme in controls and *Shirks* was quantified over 3-5 replicates at E10.5, and E11.5. Average counts were analyzed for significance using a Student’s t-test, and error bars denote the standard error of mean. *p<0.05 **p<0.005 ***p<0.0005; n.s.-not significant. All scale bars denote 50pm.

**Figure 4-figure supplement 4.**
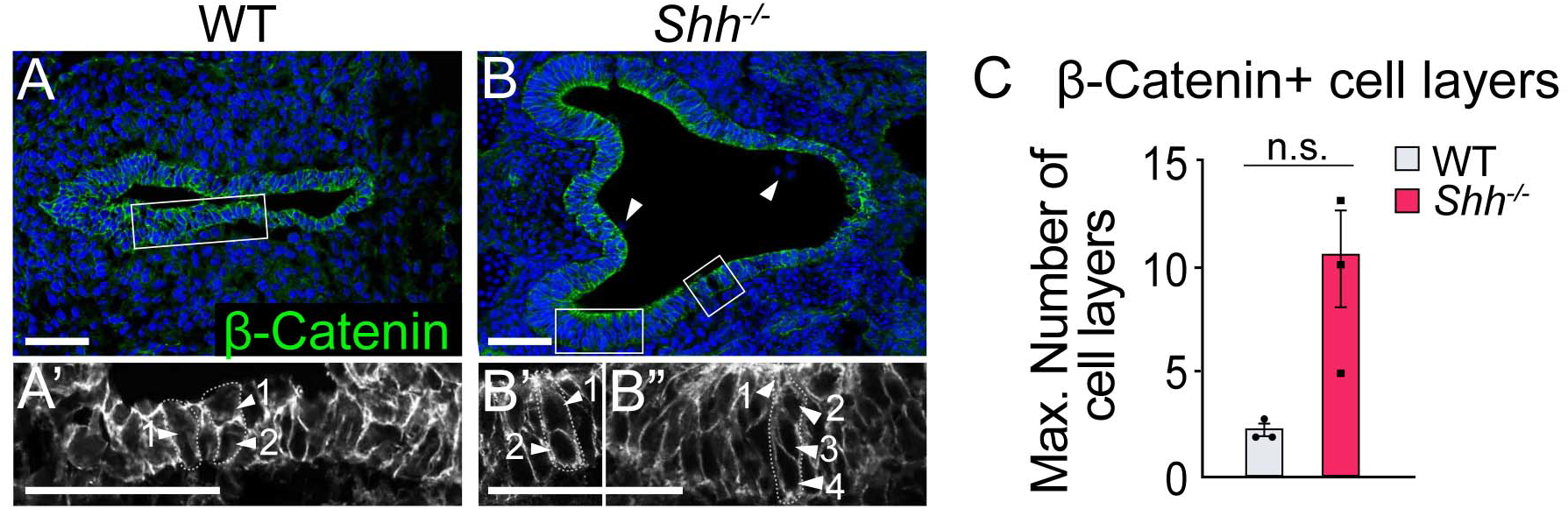
Loss of *Shh* results in a thickening and cell disorganization of the epithelial layer. A-C. 3-Catenin expression (green) marks the cell layers within the epithelium. C. The number of cell layers was averaged across 3 controls and 3 *Shh+s* and analyzed using a Student’s t-test (n.s.-not significant). Error bars show the standard error of the mean. B. Arrowheads mark extruded cells within the lumen. B’-B”. Arrowheads mark cell layers within two regions of the epithelium. All scale bars denote 50pm.

**Figure 5-figure supplement 1.**
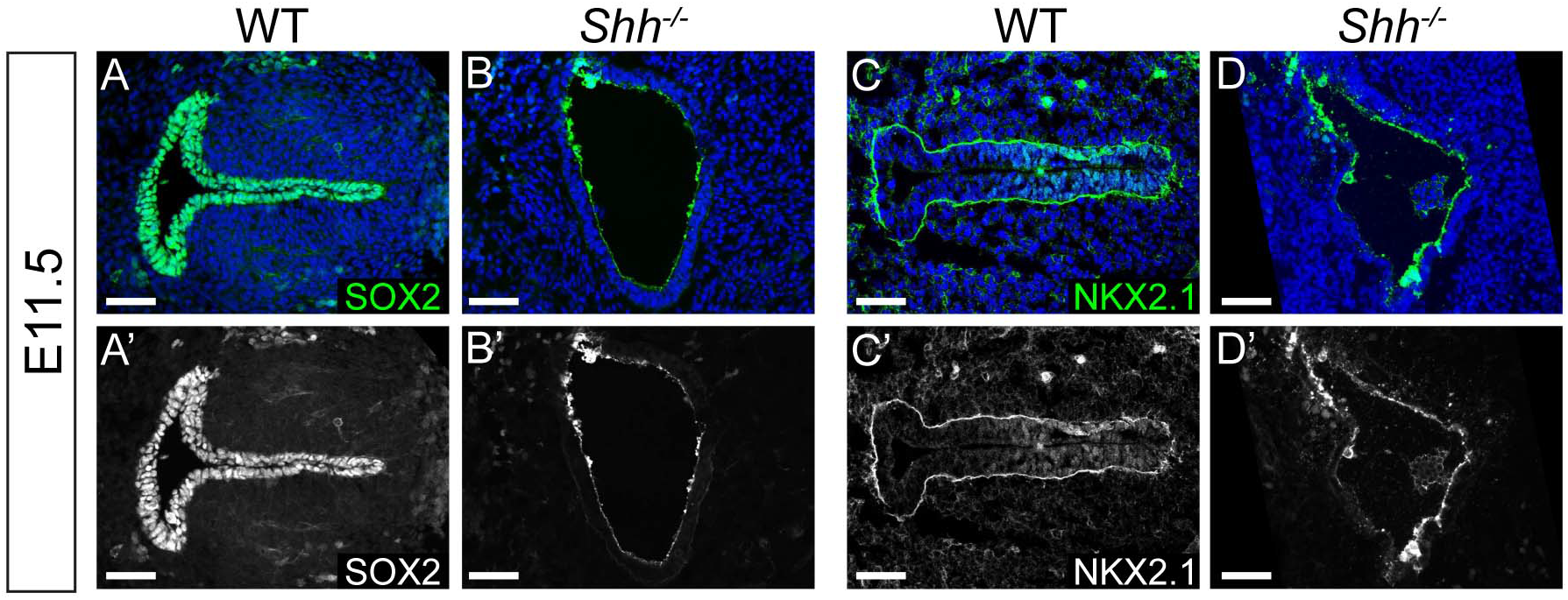
SOX2, and NKX2.1 are absent from the larynx epithelium by E11.5 in *Shir’s.* A-D. SOX2 (green; A-B) and NKX2.1 (green; C-D) expression within the epithelium of 3 control and 3 *Shir’s* at E11.5. All scale bars denote 50pm.

**Figure 5-figure supplement 2.**
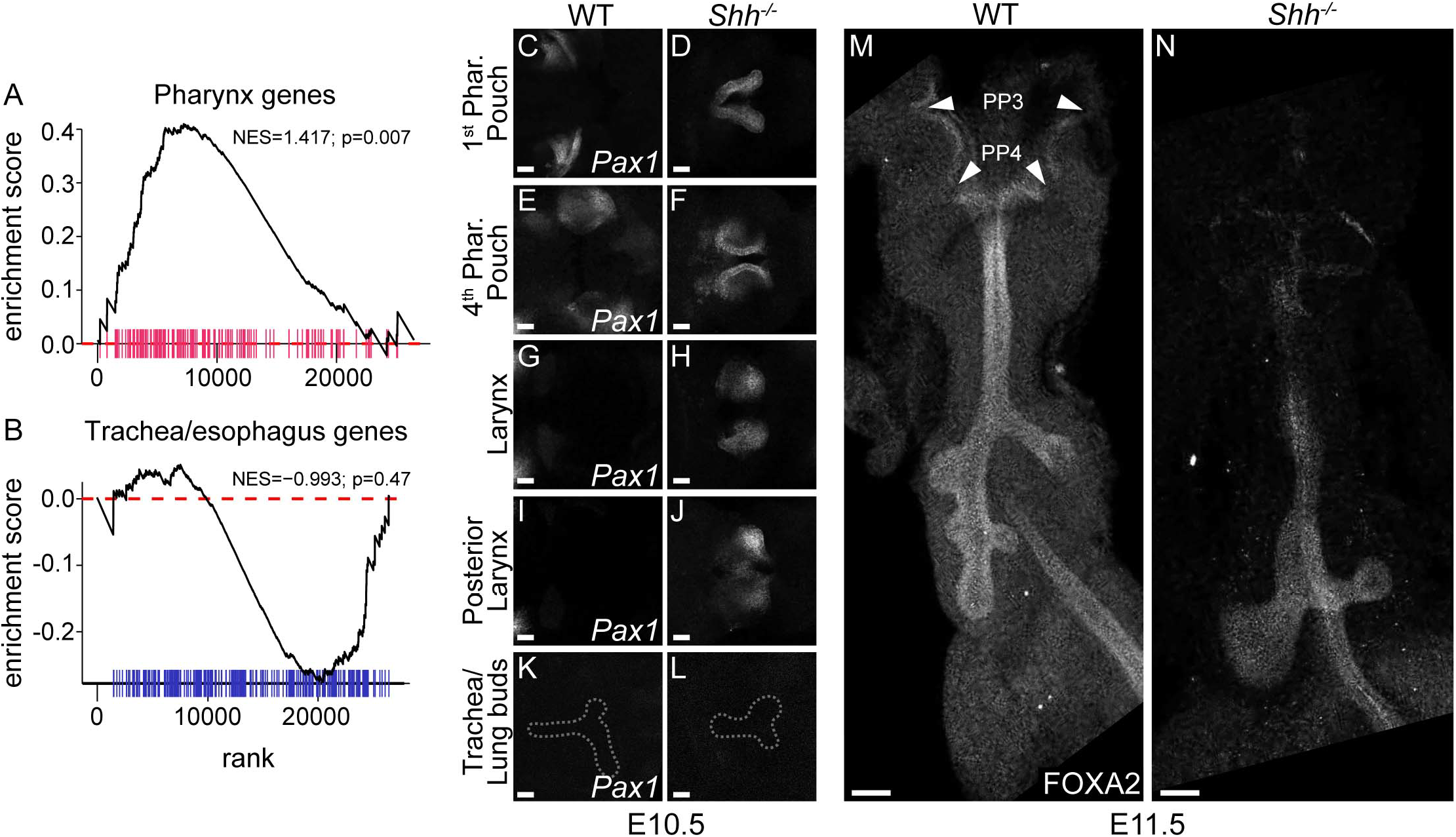
Pharynx genes are enriched while F0XA2 is lost from the anterior foregut in the absence of HH signaling. A-B. Gene-set enrichment analysis on all differentially expressed genes in *Shlr^-/-^* larynxes (Supplemental Data Table 1) against E9.5 pharynx (A, pink) and tracheal/esophageal (B, blue) endodermal genes identified by Han et al., 2020. Graphs were generated using the fgsea R package and p-values were obtained using a permutation test. C-L. Wholemount *Pax1* expression along the anterior-posterior axis of foregut epithelium in 3 control and 3 *Shir^-/-^* embryos at E10.5. Images were taken along a transverse plane. M-N. Sagittal view of wholemount FOXA2 expression along the anterior foregut in E11.5 control (3 replicates) and *Shh^-/-^* (3 replicates) embryos. NES-normalized enrichment score; PP3 and PP4 denote the 3rd and 4th pharyngeal pouches. All scale bars denote 50pm.

